# Enhancing the iNKT cell immunotherapy platform by combining optimised CAR endodomains with novel iNKT engagers

**DOI:** 10.1101/2025.10.23.683869

**Authors:** Kanagaraju Ponnusamy, Klesti Karaxhuku, Yuchao Jiang, Lyra Randzavola, Hongwei Ren, Ilia Leontari, Bryan Lye, Mehmood Zaidi, Edward J Bartlett, Edward W Tate, Vasileios Pardalis, Dimitrios Leonardos, Reza Nadafi, Irene Sarkar, Rogier M Reijmers, Marco Bua, Maria Atta, Alexia Katsarou, Irene AG Roberts, Aristeidis Chaidos, Anastasios Karadimitris

## Abstract

iNKT cells are emerging as a highly promising immunotherapy platform for the treatment of cancer. To maximise the anti-cancer activity of CAR-iNKT against the blood cancer multiple myeloma we investigated optimal CAR designs and their combination with novel iNKT-specific engagers. We find that amongst five different CAR endodomains, underpinned by increased avidity and a cross talk between Plexin D1 on CAR-iNKT and Semaphorin 4A on myeloma cells, BCMA CD28z CAR-iNKT exert the highest anti-myeloma activity. Notably, CD28z CAR-iNKT outperform their CAR-T counterparts.

To expand the anti-myeloma potential of CAR-iNKT, we designed and validated a high efficacy BCMA iNKT-specific engager which exerts significant anti-myeloma activity in conjunction with adoptively transferred iNKT cells. Finally, combined, dual target therapy with FCRL5 CAR-iNKT and BCMA iNKT engagers outperforms FCRL5 CAR-iNKT and limits immune escape of FCRL5-negative myeloma. Thus, optimised iNKT-based, dual-target, dual-modality immunotherapy has enhanced anti-tumor activity against multiple myeloma and potentially other malignancies.

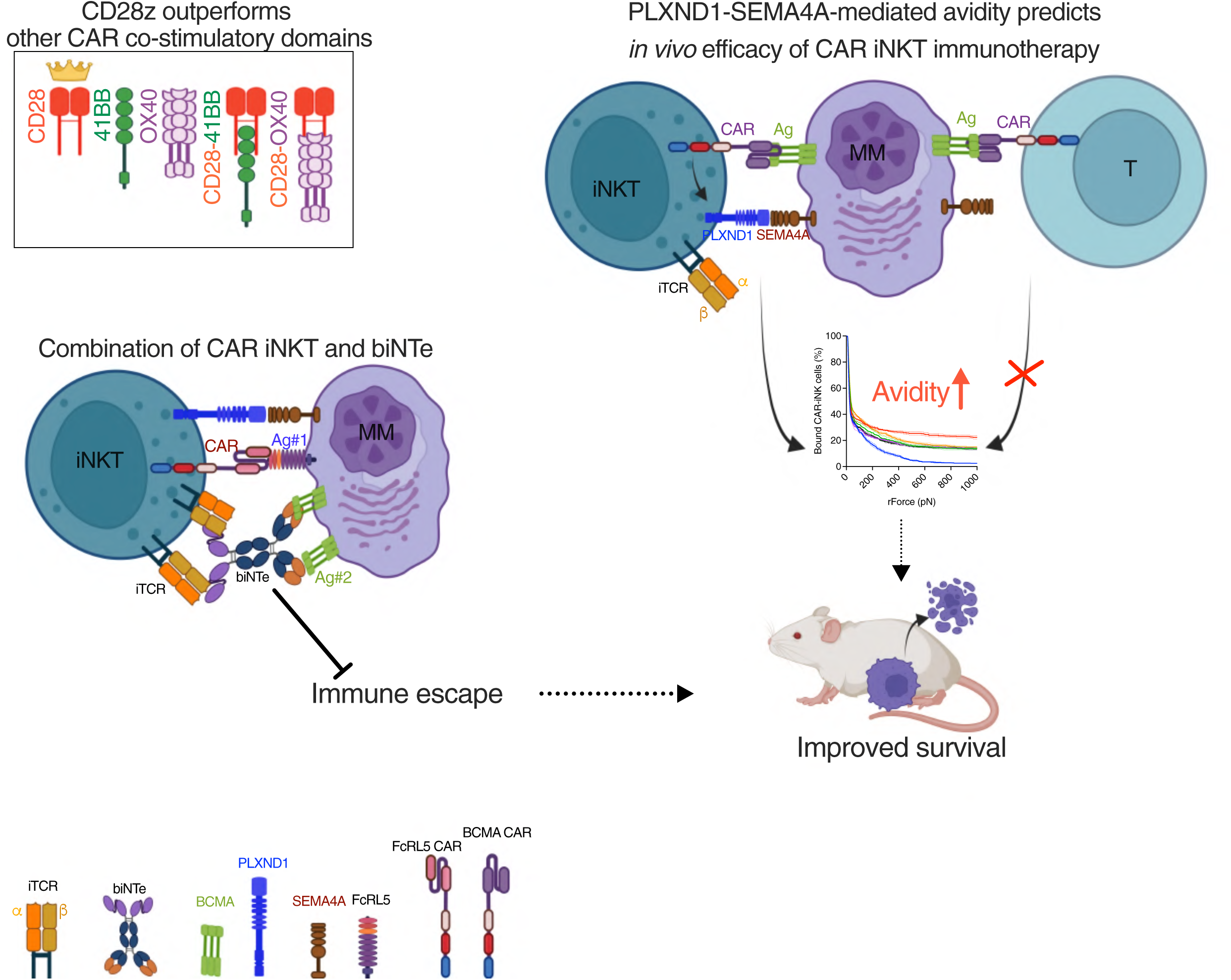

---

Cellular immunotherapies such as autologous chimeric antigen receptor (CAR)-T cells have dramatically improved the outcome of patients with some blood cancers and are potentially curative for relapsed/refractory B non-Hodgkin lymphoma^1^ and B acute lymphoblastic leukemia^2^. Cure of the B lineage malignancy multiple myeloma (MM) has proved much more difficult to achieve. However, recent results using Cilta-cel, an autologous anti-BCMA CAR-T product, showed a 15% 5-year minimal residual disease-negative state and disease-free survival in multiply relapsed disease, suggesting curative potential for some patients^3^. We hypothesise that cure rates for MM could be increased by employing a strategy that would harness the properties of effectors more powerful than T cells such as iNKT cells; optimise CARs, in particular their endodomain; and develop additional therapeutic modules such as iNKT-specific engagers targeting additional myeloma-associated antigens to allow broader targeting of myeloma plasma cells.

Given their ability to protect from acute graft-versus-host disease (aGVHD)^4,5^, iNKT are emerging as an attractive allogeneic immune cell effector platform to autologous or allogeneic T cell-based products for the treatment of blood cancers and autoimmune diseases. iNKT, a rare subset of T cells, have features of both innate and adaptive immunity and possess effector as well as immunoregulatory activity. They are characterised by an invariant TCRVα24Jα18 pairing with a TCRVβ11 chain^6^, and are restricted by the glycolipid-presenting, MHC-like molecule CD1d^7^. Their anti-tumor activity is mediated by CD1d-dependent and -independent mechanisms, including direct killing of tumor cells, antigen-presenting cell (APC) maturation and activation of tumor-specific T cells^8,9^ and depletion of immunosuppressive, CD1d-expressing tumor-associated myeloid cells^10,11^. Moreover, iNKT can be deployed as a speedier, ‘off-the-shelf’ immunotherapy platform without the need for deletion of the endogenous TCR and the additional genetic engineering this entails in allogeneic CAR-T cells, thereby reducing costs and manufacturing complexity. Additional potential advantages of allogeneic iNKT cell-over T cell-based immunotherapies include their ability to directly target CD1d-expessing tumors such as myeloma plasma cells^12^ through the iTCR; the higher potency of CAR-iNKT over CAR-T as we and others have shown^13–17^, and a reduced risk for cytokine release syndrome and neurotoxicity^18^.

Efficacy of CARs is determined by several structural features including by the choice of the endodomain such as CD28z vs 4-1BBz. CD28z CARs achieve swifter and more robust activation of T cells while 4-1BB CARs promote longer persistence of CAR-T but often fail to tackle CAR antigen low disease^19^. Here we describe a strategy to improve ‘off-the-shelf’ cellular immunotherapy for MM by engineering CAR-iNKT cells equipped with an optimal CAR endodomain that includes the critical co-stimulatory domain(s). After comparing five co-stimulatory domains we find that CD28z elicits the highest avidity against MM cells. Reflecting their enhanced avidity, CD28z CAR-iNKT outperform all other CAR-iNKT as well as CAR-T counterparts in *in vivo* models of MM. The increased avidity and *in vivo* anti-myeloma activity of CD28z CAR-iNKT is underpinned by a novel mechanism that involves interaction of PLXND1 with SEMA4A, a PLXND1 ligand highly expressed on myeloma cells. To further enhance the therapeutic potential of CD28z CAR-iNKT, we developed novel iNKT cell-specific engagers which in combination with optimized CAR-iNKT achieve broader myeloma antigen targeting and higher efficacy than single CAR-iNKT or iNKT plus bispecific engager and limit CAR target-negative disease.

## Results

### Proliferative and avidity advantage of CD28z CAR-iNKT

We generated five different lentiviral CARs against the established anti-myeloma target BCMA using the previously described and clinically validated anti-BCMA C11d5.3 scFv^20^, a G4S linker, CD8a-derived spacer and transmembranous domains and either 2^nd^ generation CD28z, 4-1BBz, OX40z or third generation CD28-4-1BBz and CD28-OX40z endodomains **(Figure 1A)**. Using our previously published protocol we then generated corresponding, same donor highly pure CAR-iNKT with similarly high CAR expression levels **(Figure 1B&C)**.

**Figure 1.**
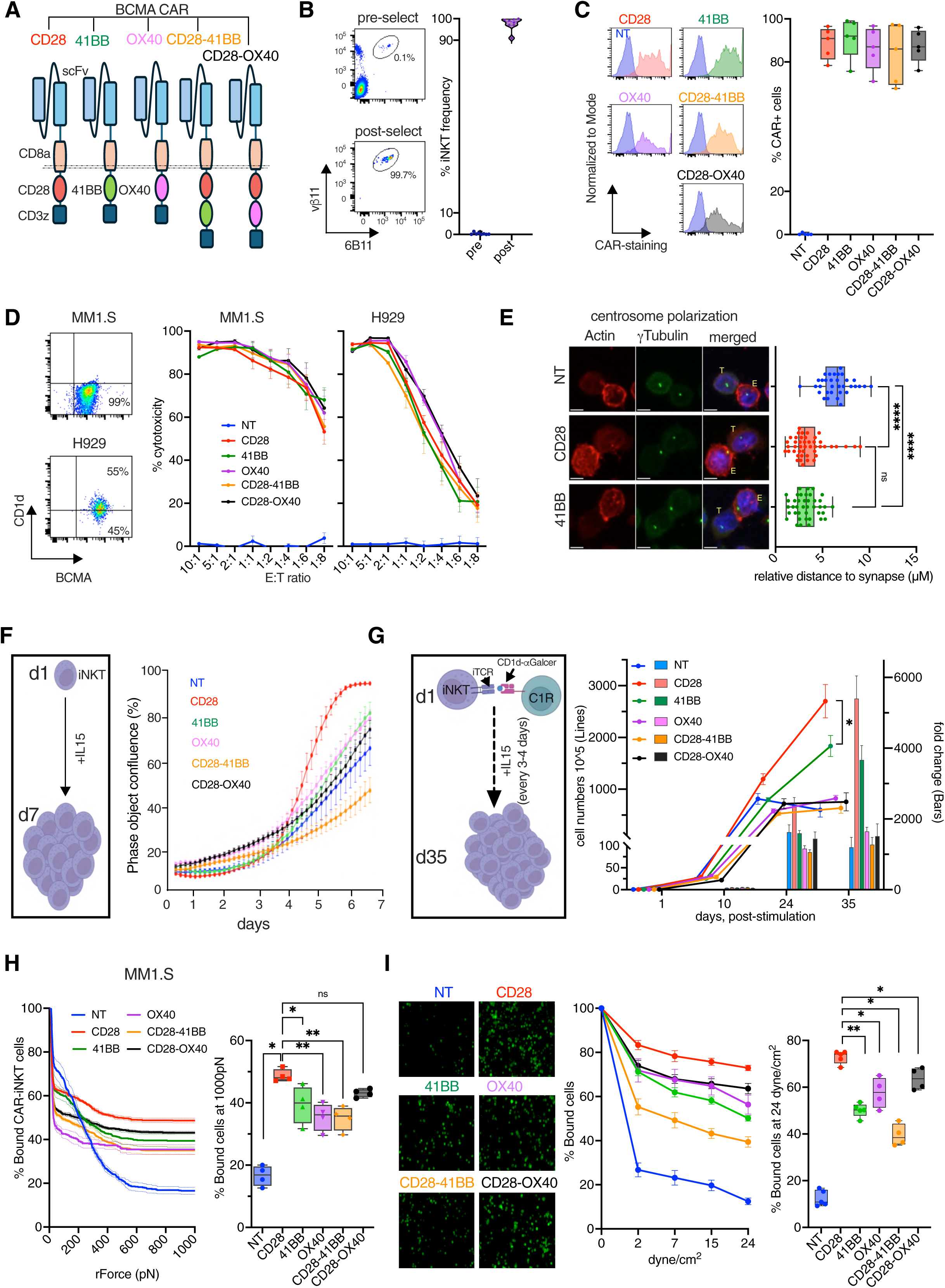
Higher proliferative potential and avidity of CD28z CAR-iNKT **A.** Structure of CARs with different co-stimulatory molecules sharing the same BCMA scFV-CD8spacer-CD8aTM design. **B.** Left: iNKT frequency in PBMC before and after iNKT microbead-based selection as identified after staining with the shown Abs and flow-cytometry. Right: cumulative data from eight different healthy donors. **C.** High transduction efficiency for all 5 CARs is shown in flow-cytometry histogram (left) and cumulative data for different donors (right; n=5). CAR staining was performed with anti-linker mAb. **D.** Surface expression of BCMA and CD1d detected by flow cytometry in MM1.S and H929 cells (left) and cytotoxic activity of BCMA CAR iNKT cells with different co-stimulatory molecules against MM1.S and H929 cells as compared to non-transduced (NT) iNKT cells are shown (representative of n=5 different donors). **E.** Immune synapse formation of BCMA CAR iNKT cells conjugated to MM1.S target cells. 3D reconstruction of z-stacks is shown and used to quantitate the distance of the centrosome (in green) to the synapse; distance < 3 µM = polarised centrosome (n=replicates of 3 different donors). **F.** Proliferative capacity of CD28z CAR-iNKT cells as compared to all other CAR- and NT iNKT cells in short-term assay in the presence of IL15 alone assessed by real-time IncuCyte®imaging (n=3 donors). **G.** Absolute numbers and fold expansion of CD28z CAR iNKT cells after stimulation with alphaGalCer-loaded C1R-CD1d cells as compared to all other CAR iNKT cells (n=3 donors). **H.** Left: CD28z CAR iNKT cell binding capacity on MM1.S myeloma cells under increasing levels of acoustic force as compared to all other CARs and NT iNKT cells; right: CAR-iNKT binding at rForce 1000pN (data representative of n=2 donors, datapoint shown for individual run). **I.** Avidity of CAR-iNKT cells against MM1.S myeloma cells as assessed under shear stress by flow chamber assay. From left: Representative image of cell binding at 24 dyne/cm^2^, cell binding efficiency at different shear tress levels, cumulative data for cell binding efficiency at 24dyne/cm^2^ (data representative of n=4 donors). ns-not significant, * p<0.05, ** p<0.01, **** p< 0.0001.

Using different donors we found that all five CAR-iNKT constructs were similarly cytotoxic against the BCMA high- and low-expressing H929 (CD1d+) and MM1.S (CD1d-) myeloma cells respectively **(Figure 1D)**. As expected, reactivity was not observed against BCMA knock out H929 or BCMA-negative K562 cell lines **(Suppl Fig. 1A&B)** confirming their specificity. Similarly, there were no clear differences in production of granzyme B and interferon-γ **(Suppl Fig. 1C&D)** and, although confocal microscopy imaging showed all CAR-iNKT established more pronounced immune synapses than non-tranduced (NT) iNKT cells with myeloma cells as assessed by centrosome polarisation at the immune synapse, again there were no differences between them **(Figure 1E and Suppl Fig. 1E).**

However, short term proliferation assays performed by real time imaging **(Figure 1F)** as well as assessment of long-term expansion potential in the presence of IL-15 and re-stimulation with CD1d-expressing B cell lines loaded with the glycolipid alpha-galactosylceramide **(Figure 1G)**, showed that CD28z CAR-iNKT were significantly more proliferative than CAR-iNKT with the other four endodomains including 4-1BBz CAR-iNKT.

A more detailed functional analysis focused on CD28z vs 4-1BBz CAR-iNKT showed that they had the same cytotoxic activity against an extended panel of myeloma cell targets **(Suppl Fig. 1F-H)** independent of the CAR expression levels **(Suppl Fig. 1I-J)**.

As *in vivo* potency of CAR-T cells is often better predicted by their avidity against their tumor targets^17,21–23^, we employed both acoustic force- and shear stress-based avidity assays **(Figure 1H&I)** to study the strength of CAR-iNKT cell interaction with myeloma cells at steady state. We found that in both assays and using two different myeloma cells as targets, CD28z CAR-iNKT demonstrated the highest avidity amongst all five CAR-iNKT designs **(Figure 1H&I and Suppl Fig. 1K-L)**.

Therefore, higher proliferative potential and avidity distinguish CD28z from other 2^nd^ and 3^rd^ generation endodomains in the context of CAR-iNKT-mediated targeting of MM.

### PLXND1-SEMA4A interaction determines higher avidity of CD28z CAR-iNKT

Next, we investigated the mechanism that underpins higher avidity of CD28z CAR-iNKT. Since avidity is measured within minutes of adding resting effectors to their target cells we reasoned that differences would be mediated by proteins already expressed on the surface of the different types of CAR-iNKT cells. To identify such proteins, we compared the transcriptomes by RNA-sequencing of resting CD28z and 4-1BBz CAR-iNKT as well as NT NKT from three different donors. We found a small number of differentially expressed genes between CD28z or 4-1BBz CAR-iNKT vs iNKT cells and 28z vs 4-1BBz CAR-iNKT **(Figure 2A-C and Suppl Table 1).** Both CD28z and 4-1BBz CAR-iNKT upregulate key genes related to cell adhesion and migration such as *ITGAD*, *MMP9*, *COL6A2* and also *IFNG* **(Figure 2C)**. Amongst the few differentially expressed genes, we found that *PLXND1*, a semaphorin ligand family receptor was significantly upregulated in CD28z CAR-iNKT **(Figure 2B&C)**. Flow cytometry analysis confirmed significantly higher levels of expression of PLXND1 in CD28z CAR-iNKT as compared to 4-1BBz, other CAR-iNKT designs and NT iNKT **(Figure 2D and Suppl Fig. 2A)**, a finding also corroborated by quantitative immunofluorescence **(Figure 2E)**. Expression of PLXND1 remained stable and did not increase upon CAR-iNKT activation in the presence of myeloma cells **(Suppl Fig. 2B)** while no significant differences of PLXND1 expression between CD4+ and CD4-iNKT cells were observed **(Suppl Fig. 2C)**. In contrast to CAR-iNKT, PLXND1 was not detected on CD28z or 4-1BBz CAR-T cells by flow-cytometric analysis **(Figure 2D)**. In support of this, published single cell RNA-seq data shows low level *PLXND1* expression in memory T cells but no expression in naive T or T-reg cells **(Suppl Fig. 2D)**.

**Figure 2.**
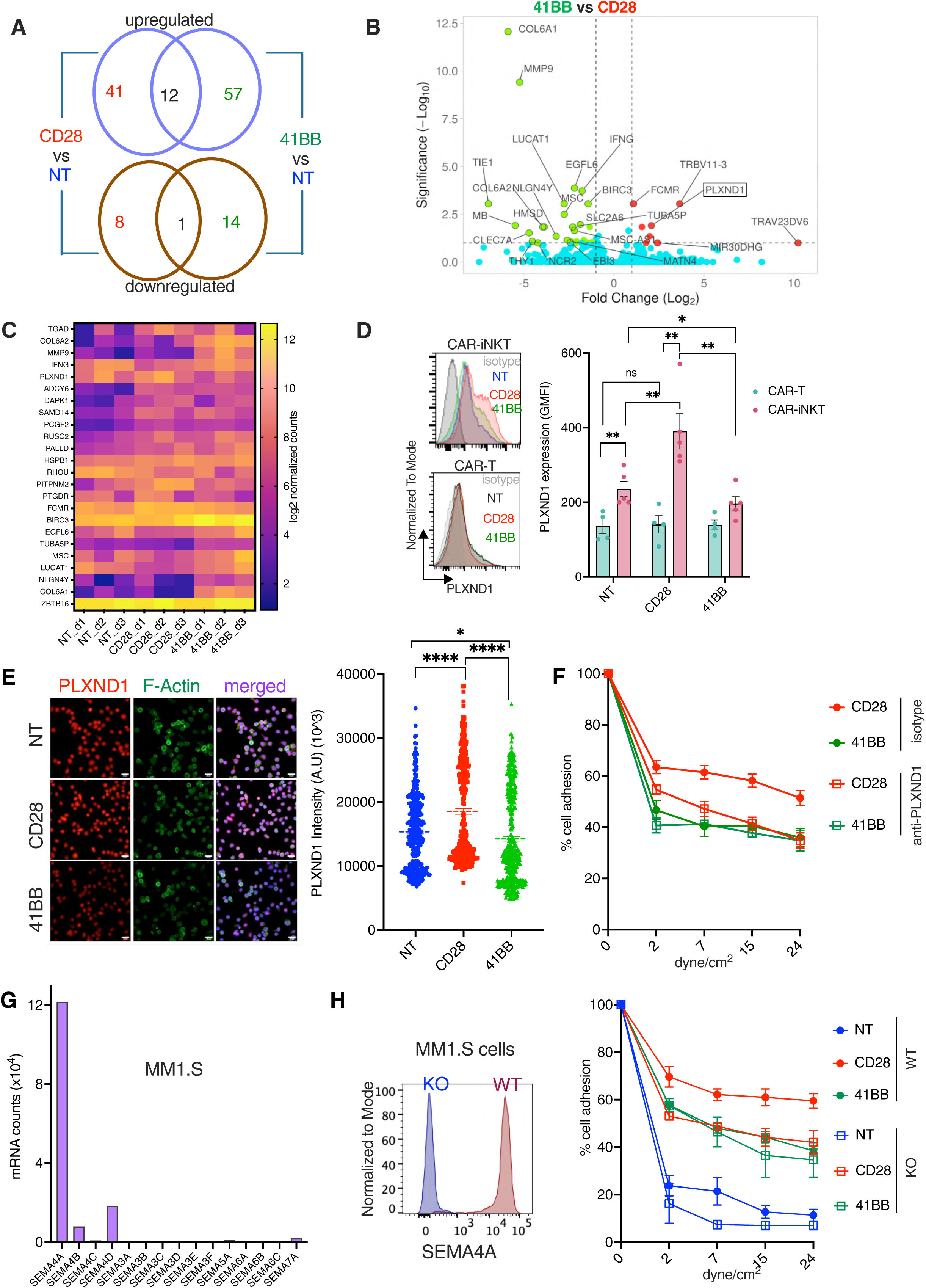
PLXND1-SEMA4A interaction determines higher avidity of CD28z CAR-iNKT **A.** Venn diagram shows number of commonly shared differentially regulated genes between CD28z and 41BBz compared to NT iNKT cells at steady state (n=3 donors (d1-d3)). **B.** Volcano plot for differentially regulated genes between CD28z and 41BBz CAR iNKT cells at steady state (n=3 donors). The names of top 25 significantly regulated genes are shown including PLXND1. **C.** Heatmap shows the log2 count values of key genes differentially expressed in NT, CD28z and 41BBz CAR iNKT cells. ZBTB16, a marker for NKT cells shown as control. (n=3 donors). **D.** PLXND1 expression in CD28z- and 4-1BBz-CAR iNKT cells as compared to NT iNKT as assessed by flow-cytometry (histogram, top) and in T cells (histogram, bottom); right: Geometric Mean of Fluorescence Intensity (GMFI) cumulative data (n=4-5 donors). **E.** Left: PLXND1 expression in CD28z CAR iNKT cells as compared to NT and 4-1BBz iNKT cells as assessed by immunofluorescence; right: cumulative data of intensity of staining. Each dot represents a cell (n=3 pooled donors). **F.** Cell binding efficiency of CD28z CAR iNKT cells on MM1.S cells in the presence of PLXND1-blocking antibody or isotype control under shear stress (n=4 donors). **G.** Expression of SEMA4A mRNA and its family members in MM1.S cells from CCLE dataset. **H.** Left: SEMA4A expression in SEMA4A in wild type and knock-out MM1.S cells. right: Avidity of NT, CD28 and 41BB CAR-iNKT cells under shear stress against SEMA4A-expressing and non-expressing MM1.S cells (n=4 donors). ns-not significant, * p<0.05, ** p<0.01, **** p< 0.0001.

To further explore the role of PLXND1 we repeated the shear stress avidity assays in the presence of a blocking PLXND1 mAb^24^. In relation to Ig isotype control, this resulted in a dose-dependent reduction of CD28z CAR-iNKT avidity but had no effect on the avidity status of 4-1BBz counterparts **(Figure 2F and Suppl Fig. 2E)**.

Therefore, the higher avidity of CD28z CAR-iNKT over NT and 4-1BBz CAR-iNKT is PLXND1-dependent.

To investigate this further we measured gene expression of *SEMA4A* and *SEMA3E*, the semaphorin family members which could potentially act as ligands for PLXND1^25–27^, in the two myeloma cell lines used in the avidity assays. While expression of *SEMA4A* was the highest amongst all semaphorins, *SEMA3E* was not expressed in either MM1.S or H929 cells **(Figure 2G and Suppl Fig. 2F)** with the same expression pattern seen in primary myeloma plasma cells from patients with MM **(Suppl Fig. 2G)**. Of note, SEMA4A has been validated as a therapeutic target for CAR-and Ab-based immunotherapy of MM and similar to FCRL5^28,29^, another clinically validated bispecific engager target, its expression is highest in the high risk chr1q-amp myeloma cells^30^ **(Suppl Fig. 2H)**.

To confirm the role of SEMA4A in CAR-iNKT avidity, using CRISPR/Cas9 gene editing we generated MM1.S myeloma cells lacking expression of SEMA4A **(Figure 2H).** Although the cytotoxic activity of CD28z and 4-1BBz CAR-iNKT was the same against both SEMA4A-expressing and gene-edited SEMA4A-negative MM1.S cells **(Suppl Fig. 2I)**, avidity of CD28z CAR-iNKT against SEMA4A-negative myeloma cells was lower compared to SEMA4A-expressing myeloma cells, and at the same level as that of the 4-1BBz CAR-iNKT cells which was unaffected by abolishing SEMA4A expression **(Figure 2H).** These findings demonstrate that enhanced avidity of CD28z CAR-iNKT requires expression of SEMA4A on myeloma cells.

Finally, we tested whether differential cell proliferation and avidity induced by CD28z co-stimulatory domain would be independent of the CAR target. For this purpose, we synthesised 4-1BBz and CD28z CARs against FCRL5. The scFv of these CARs was based on the clinically active FCRL5 engager cevostomab^30^ and was previously validated in the context of CAR-T therapy^31^. We found that as for BCMA CAR-iNKT, CD28z and 4-1BBz FCRL5 CAR-iNKT cells show similar cytotoxicity against FCRL5-expressing MM1.S myeloma cells **(Figure 6A and Suppl Fig. 2J)**. However, CD28z FcRL5 CAR iNKT cells showed higher proliferative potential than 4-1BBz FcRL5 CAR iNKT or NT iNKT cells **(Suppl Fig. 2K)** and higher expression of PLXND1 than their 41BBz counterparts commensurate with higher PLXND1-dependent avidity against FcRL5+ MM1.S cells **(Suppl Fig. 2L)**.

Together these findings demonstrate that the higher avidity of CD28z CAR-iNKT is mediated by its higher expression of PLXND1 and its ability to engage SEMA4A on myeloma cells and it is independent of the CAR scFv and its myeloma target.

### CD28z CAR-iNKT exert a more effective anti-myeloma activity *in vivo* in a PLXND1-dependent manner

To investigate whether avidity indeed predicts anti-myeloma activity *in vivo*, we employed a systemic model of MM using luciferase- and tdTomato-expressing MM1.S (MM1.S-Luc) myeloma cells and monitored tumor burden by bioluminescence imaging (BLI). To better delineate differences between CAR constructs we used the limiting dose^32^ of 10^6^ CAR-iNKT cells per animal for each group **(Figure 3A)**. We found that CD28z CAR-iNKT followed by CD28-OX40z CAR-iNKT were the most potent in delaying tumor growth amongst the five CAR types as assessed by BLI **(Figure 3B&C)** while prolongation of survival was more pronounced in animals treated with CD28z CAR iNKT **(Figure 3D)**.

**Figure 3.**
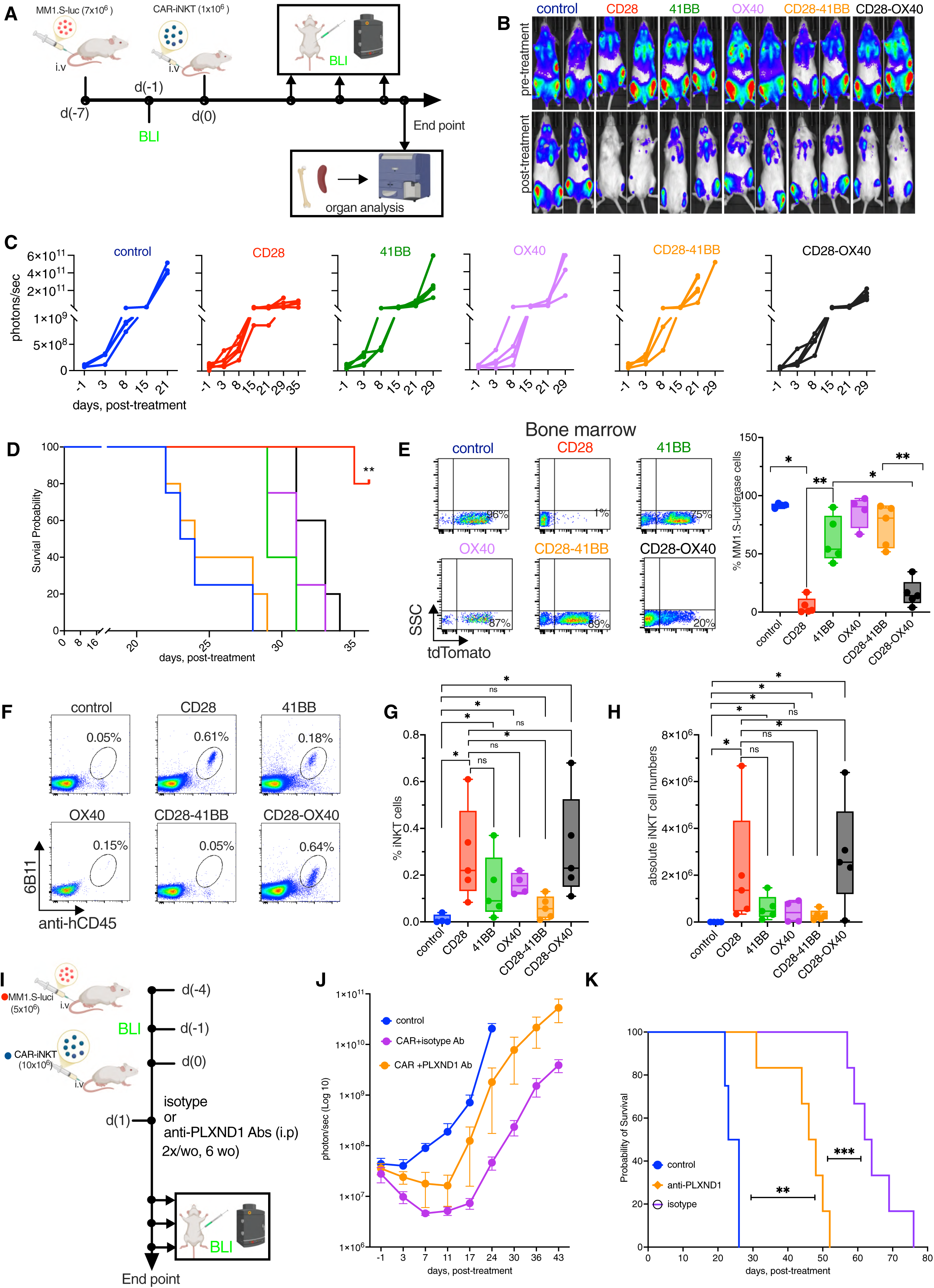
CD28z CAR-iNKT outperform other CAR designs in PLXND1-dependent manner **A.** Schematic of *in vivo* experimental design associated with figure 3 (A-H). **B-C.** Representative BLI images of mice from control and BCMA CAR iNKT with different co-stimulatory molecules-treated groups for before treatment and post-treatment (week 3) and its cumulative photon strength shown for each group (n=4-5 mice per group). **D.** Kaplan-Meier survival curve for all the groups treated with 1×10^6^ BCMA CAR iNKT cells and control group (n=4-5 mice per group). **E.** Left: Representative flow cytometry dot plots and its cumulative data (right) for frequency of MM1.S-luci cells as measured by tdTomato fluorescence marker by flow cytometry in the bone marrow of all the groups collected at end point (n=4-5 mice per group). **F&G.** Representative flow cytometry analysis plots for iNKT frequency in the bone marrow for each group (F), and its cumulative data (G). **H.** Absolute iNKT cell numbers in the bone marrow of each group counted based on flow cytometry analysis using counting beads. **I.** Schematic of *in vivo* experimental design associated with figure 3J&K. **J&K.** Total body BLI signals of mice treated with anti-Plexin D1 antibody treated group as compared to isotype antibody-treated group and untreated control group (J) and its Kaplan-Meier survival curve (K) (n=4-6 mice per group). ns-not significant, * p<0.05, ** p<0.01.

Immunophenotypic analysis of bone marrow, spleen and peripheral blood confirmed significantly lower myeloma burden in CD28z CAR iNKT-treated mice **(Figure 3E and Suppl Fig. 3A&B)**. The CAR-iNKT frequency and absolute numbers in the bone marrow at end point, although did not reach statistical significance, were higher in CD28z and CD28-OX40 CAR-iNKT compared to other CAR-iNKT groups **(Figure 3F-H)**. Together these data validate the avidity assay prediction that CD28z CAR-iNKT are the most potent amongst the five CAR-iNKT groups.

Finally, we investigated the impact of PLXND1 blockade *in vivo* by treating myeloma-bearing mice with CD28z CAR-iNKT cells plus either PLXND1 blocking Ab or isotype (Ig) control as shown in **(Figure 3I)**. While both groups of CAR iNKT were effective in controlling disease progression as compared to untreated controls, survival of CAR-iNKT plus PLXND1 Ab-treated animals was significantly shorter than that of CAR-iNKT plus Ig-treated animals **(Figure 3J&K)**. Therefore, we conclude that PLXND1-dependent higher avidity accounts for the higher anti-myeloma activity of CD28z CAR-iNKT *in vivo*.

### CAR-iNKT outperform CAR-T against myeloma

Next, since our previous work demonstrated that iNKT cells are a more powerful immunotherapy platform for CAR therapy of lymphoma and high risk ALL^13,17^, we investigated whether this is also the case in MM. Indeed, CD28z BCMA CAR-iNKT from three different donors were consistently more cytotoxic towards MM1.S myeloma cells than their CAR-T counterparts **(Suppl Fig. 4A)** and exhibited higher PLXND1-dependent avidity *in vitro* than CAR-T **(Figure 4A)**. In line with these observations when used at the limiting dose of 10^6^ CAR+ effectors **(Figure 4B)**, CAR-iNKT were significantly more potent than CAR-T counterparts *in vivo* in delaying growth of systemic myeloma **(Figure 4C&D)** and depleting myeloma cells from bone marrow and spleen **(Figure 4E&F)**. Accordingly, survival of CAR-iNKT-treated mice was significantly longer than that of CAR-T counterparts **(Figure 4G).**

**Figure 4.**
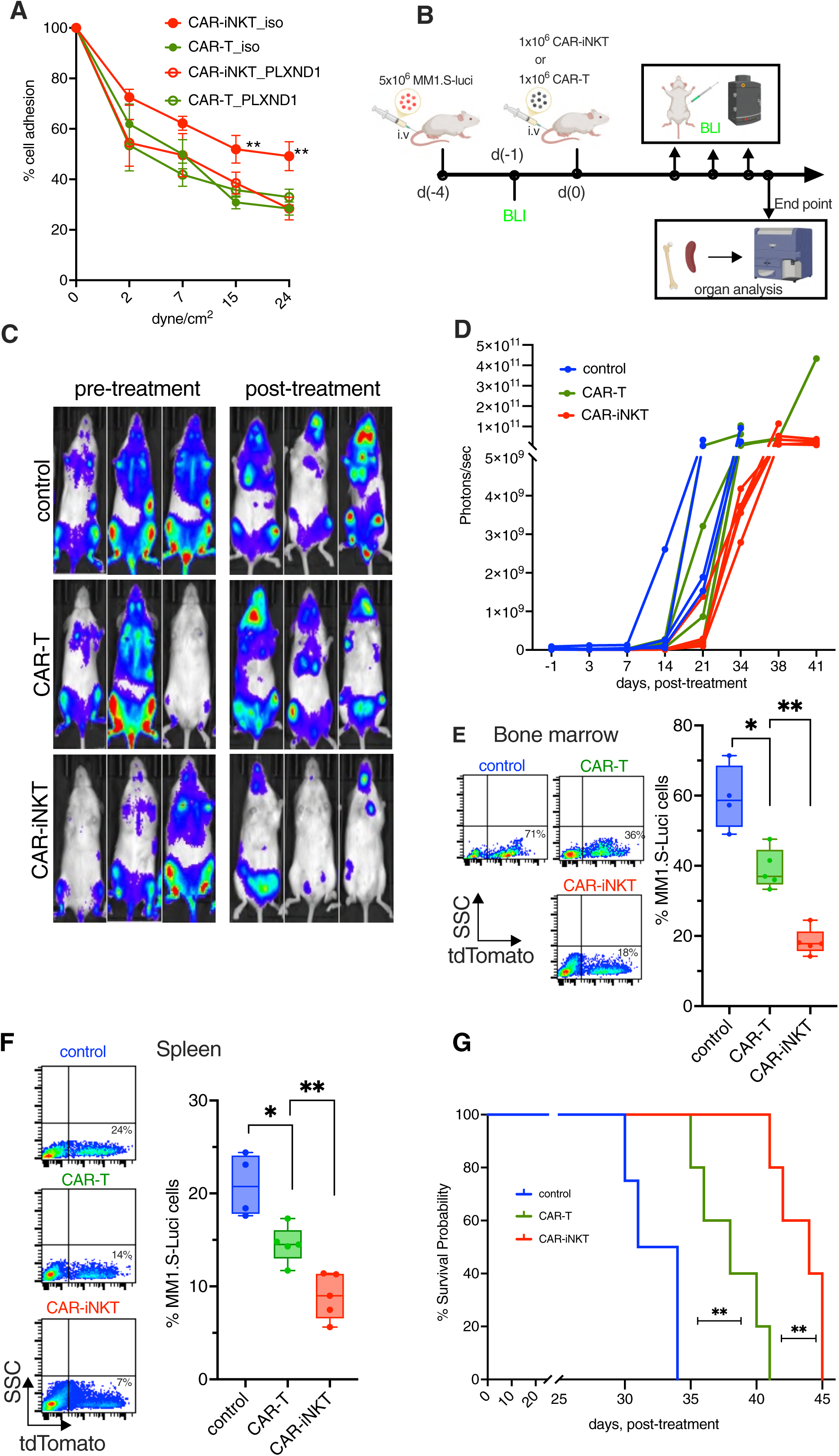
CAR-iNKT outperform CAR-T against myeloma **A.** BCMA CAR iNKT cell avidity on MM1.S cells shown as compared to BCMA CAR-T cells of same donors with isotype or PLXND1 blocking antibody at 1µg/ml under shear stress (n=3 donors). ** Significance shown for CAR iNKT isotype compared to CAR-T isotype control. **B.** Schematic of *in vivo* experimental design associated with Figure 4. **C&D.** Representative BLI images of mice from control, BCMA CAR iNKT and BCMA CAR-T cell-treated group before and after treatment (week 3) (C) and its cumulative photon emission shown for each group (D) (n=4-5 mice per group). **E&F.** Left: Representative dot plots and cumulative data (right) for frequency of MM1.S-luci cells in the bone marrow (E) and spleen (F) analysed by flow cytometry using tdTomato fluorescence marker at end point (n=4-5 mice per group). **G.** Kaplan-Meier survival curve for myeloma-bearing mice treated with CAR iNKT or CAR-T as compared to untreated control as shown in B. (n=4-5 mice per group). * p<0.05 ** p<0.01

These data show that CD28z BCMA CAR-iNKT outperform CAR-T counterparts against myeloma, likely in a PLXND1-dependent manner.

### Development, specificity and anti-myeloma activity of iNKT cell engagers

Having identified the optimal CAR endodomain, we hypothesised that CAR-iNKT-based immunotherapy could be further enhanced by developing iNKT cell-specific engagers that would target additional myeloma-associated antigens. Such engagers would bring CAR-iNKT cells to the proximity of tumor cells facilitating their killing, as previously described for T and NK cell-specific bi- and tri-specific engagers in pre-clinical/clinical development or already licensed for clinical use^33^.

To provide proof-of-concept in support of this hypothesis, we designed two bispecific iNKT engagers (biNTe). In principle, an iNKT engager could be designed to engage either the TCRVβ11 or TCRVα24 variable, non-CDR3 segments of the iTCR or the TCRVα24Jα18 CDR3 idiotype. Here, we designed two engagers against TCRVβ11, based on the TCRVβ11 Ab clone C21^6^. Since engagement of iTCR with CD1d and lipid ligands involves primarily the TCRVα24Jα18 CDR3 such an approach would be expected to allow iNKT activation through iTCR-CD1d interaction even in the presence of a TCRVβ11 engager.

The tumor targeting arm of the engagers was designed to bind to BCMA, and like the CAR, it was based on the C11d5.3 scFv^20^ with modifications in the linker.

One biNTe design comprised a heterodimeric, monovalent, scFv-Fc human wild type IgG1 with Fc-silent (L234A/L235A (LALA) and a ‘key-in-hole’ changes **(Figure 5A)**. A second biNTe design comprised a homodimeric, bivalent IgG-scFv human wild type IgG1. The anti-BCMA arm has a Fab configuration while the anti-TCRVb11 arm has a scFv configuration **(Figure 5A)**.

**Figure 5.**
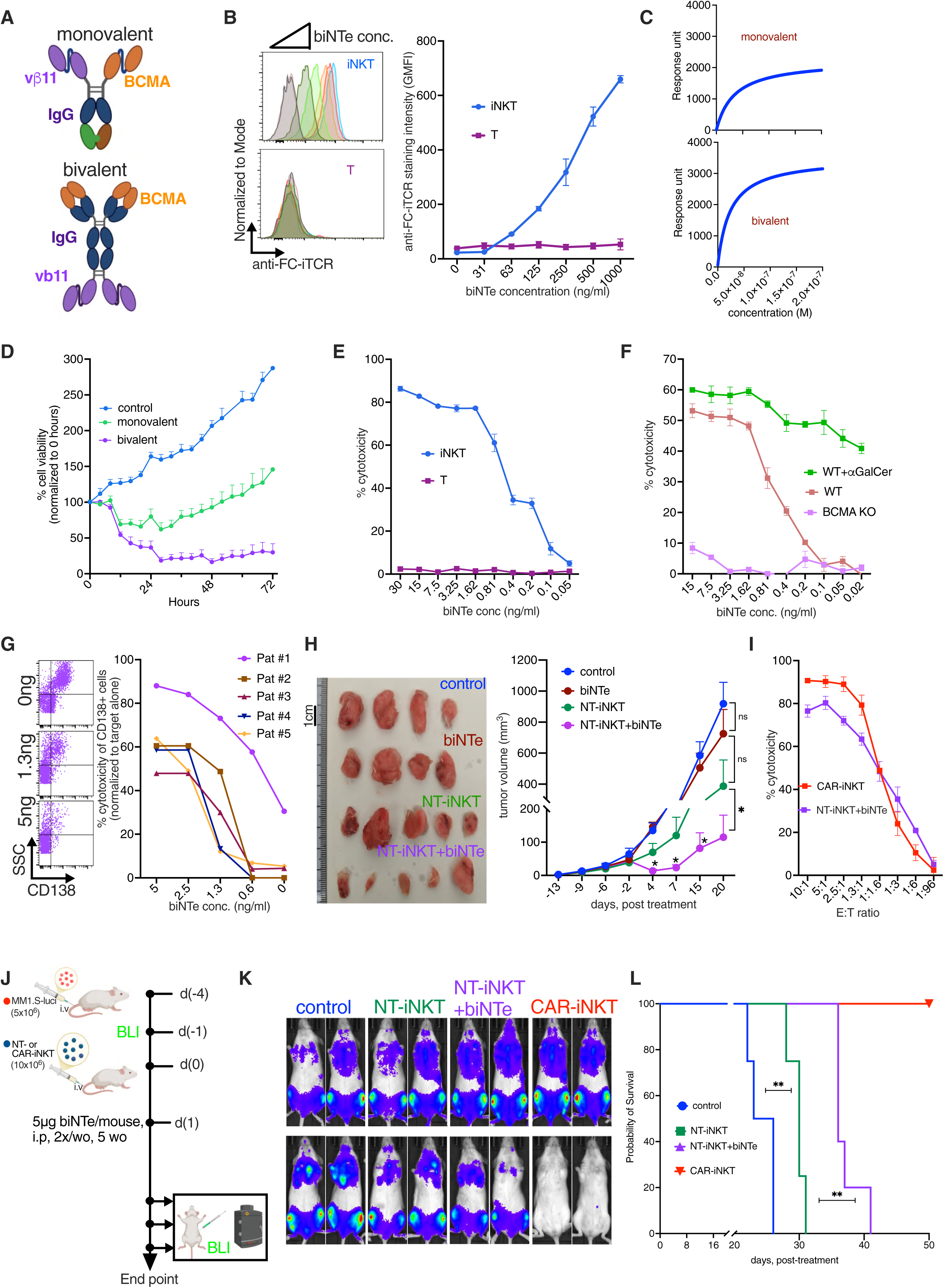
Development and anti-myeloma activity of iNKT specific engagers **A.** Design of monovalent (top) and bivalent biNTe (bottom) shown. **B.** Left: Flow cytometry analysis of same donor purified iNKT and T cells after staining with increasing concentrations (0-1µg) of bivalent biNTe and its cumulative data shown (right) as GMFI values for iNKT and T cells (n=3 donors). **C.** Representative data showing higher affinity by bivalent antibody to soluble BCMA as compared to monovalent antibody and determined by Surface Plasmon Resonance (SPR) (n=3 donors). **D.** Increased cytotoxic activity of NT iNKT cells incubated with bivalent antibody against MM1.S cells as compared to monovalent antibody as assessed by live imaging in Incucyte Zoom for up to 72 hours. MM1.S cells incubated with NT iNKT alone as control. **E.** Cytotoxic activity of same donor iNKT and T cells incubated with bivalent biNTe against MM1.S myeloma cells (representative n=3 donors). **F.** Cytotoxic activity of iNKT Incubated with biNTe in the presence or not of alphaGalCer against BCMA-expressing parental and BCMA knock out CD1d+ H929 myeloma cells. **G.** iNKT-biNTe-mediated cytotoxicity of patient-derived primary bone marrow myeloma cells at 2:1 E:T ratio and different biNTe antibody concentrations for 20hr. Left: flow-cytometric representative dot plot example of primary cells for patient #1. Right graph: cumulative data normalized to target alone (n=5 patients). CD138+ cells were measured for each condition and normalized to myeloma target cells alone. **H.** Tumor size (left) and tumor volume (right) shown from MM1.S myeloma subcutaneous tumor model treated with NT iNKT cells and/or biNTe and control mice (n=4-5 mice per group). * Significance compares the NT iNKT alone and NT iNKT+biNTe groups. **I.** Cytotoxic activity of NT iNKT cells incubated with biNTe antibody and BCMA CAR iNKT cells against MM1.S cells shown for 16hr (representative of n=3 donors). **J.** Schematic of experimental design shown for MM systemic model for biNTe treatment related to Figure 5K&L. **K.** Left: Representative BLI images of mice with systemic myeloma treated with BCMA CAR-iNKT or NT iNKT plus biNTe before and after treatment (3 weeks), and their cumulative data shown in suppl Fig 5I for (n=4-5 mice per group). **L.** Kaplan-Meier survival of myeloma-bearing mice treated as shown in Figure 5J. ns-not significant, * p<0.05, ** p<0.01.

After cloning of the constructs into an expression vector and their transfection to 293T cells, protein was purified from the culture supernatant using Protein A columns and biNTe purity was confirmed by SDS PAGE **(Suppl Fig. 5A)**.

Using the biNTe to stain purified iNKT and T cells followed by fluorescent-labelled secondary anti-human IgG1 Fc Ab, we found that compared to controls, biNTe-stained purified iNKT but not T cells in a dose-dependent manner **(Figure 5B & Suppl Fig. 5B)** thus demonstrating the specific binding of the iNKT cell arm of biNTe to the iNKT TCRVβ11 chain.

We next confirmed the ability of the tumor-binding arm of biNTe to bind their intended target BCMA showing that staining of iNKT cells occurred only in the presence of bio-sBCMA **(Suppl Fig. 5C)**. To assess the potency of binding we employed by surface plasmon resonance. This showed a lower Kd and thus higher affinity for bivalent over monovalent biNTe to soluble BCMA (Kd of 23.4nM vs 34.5nM respectively) **(Figure 5C and Suppl Fig. 5D-F)**. In line with these findings, EC50 measured against MM1.S cells in the presence of iNKT was lower for bivalent than monovalent biNTe (2.4pM vs 6.2pM respectively) **(Suppl Fig. 5G&H)**. Similarly, the bivalent biNTe demonstrated significantly higher longitudinal cytotoxic potential *in vitro* compared to the monovalent engager **(Figure 5D**). Based on its higher affinity and cytotoxic potential the bivalent engager was selected for further study.

Bivalent biNTe selectively killed myeloma cells in the presence of iNKT, but not T cells supporting the specificity of the biNTe iTCR arm **(Figure 5E)**. In addition, the anti-myeloma cytotoxic activity of iNKT against CD1d-expressing BCMA+ (but not BCMA-) H929 myeloma cells was further enhanced in the presence of biNTe and aGalCer **(Figure 5F)** confirming the BCMA specificity of the tumor engaging arm of the biNTe and BCMA- and iNKT cell-dependent killing of myeloma cells. These findings also validate the prediction that binding of the designed biNTe to the TCRVb11 chain would allow the iTCR to engage with CD1d on tumor cells and to further enhance anti-myeloma activity.

We next co-cultured bone marrow cells from five patients with myeloma with iNKT cells and varying concentrations of biNTe. We found that while the frequency of myeloma PC dramatically decreased in a biNTe dose-dependent manner, that of the non-myeloma bone marrow cells was considerably less affected, suggesting specific killing of the myeloma but not of the non-myeloma bone marrow cells **(Figure 5G and Suppl Fig. 5I).**

To corroborate the *in vitro* findings, we tested the *in vivo* efficacy of the biNTe in a subcutaneous myeloma model using the BCMA-expressing MM1.S cells. After tumor engraftment, mice were left untreated or treated with either 10^7^ unmodified, pre-expanded iNKT and 5μg/mouse biNTe ip 24hours post-iNKT treatment and then twice a week **(Figure 5H).** While iNKT or biNTe alone had no impact on tumor growth, the iNKT and biNTe combination significantly delayed myeloma tumour growth, first observed two days after the first engager dose **(Figure 5H)** thus providing proof-of-principle that biNTe when combined with adoptive iNKT immunotherapy has significant anti-myeloma activity.

We next compared the anti-myeloma activity of BCMA biNTe with that of BCMA CAR-iNKT cells. Myeloma MM1.S cells were co-cultured with NT iNKT and biNTe or BCMA CAR-iNKT at a 1:1 E:T ratio. Cytotoxicity against MM1.S cells showed slightly lower killing by biNTe plus iNKT compared to CAR iNKT attesting to the potency of the BCMA biNTe **(Figure 5I).** Finally, we compared the efficacy of BCMA biNTe plus iNKT vs BCMA CAR-iNKT in a systemic myeloma model using 10^7^ effectors **(Figure 5J).** We found that although CAR-iNKT were overall more efficacious *in vivo*, the BCMA biNTe plus iNKT combination also significantly delayed disease progression as assessed by BLI, reduced myeloma cell frequency in bone marrow and increased overall survival as compared to iNKT only-treated and untreated controls **(Figure 5K&L and Suppl Fig. 5J&K)**.

We conclude that biNTe are specific for iNKT and in combination with adoptive transfer of *in vitro* expanded iNKT exert a considerable anti-myeloma activity.

### biNTe and CAR-iNKT for dual targeting of multiple myeloma

Having established an optimal CAR endodomain for CAR-iNKT treatment of myeloma and proof-of-principle of the anti-myeloma activity of biNTe, we next tested whether combining the two therapeutic modules would enhance the anti-myeloma activity of the iNKT cell platform by targeting two different antigens simultaneously, i.e., the clinically validated targets BCMA and FCRL5 by bivalent biNTe and 28z CAR respectively.

For this purpose, we first established that FcRL5 CD28z CAR iNKT cells are cytotoxic against FCRL5+ but not FCRL5-MM1.S myeloma cells **(Figure 6A**). We then used a systemic myeloma model generated by a 50-50 mixture of BCMA+FCRL5+ and BCMA+FCRL5-MM1.S-Luc cells, i.e., BCMA and FCRL5 are expressed by 100% and 50% respectively of tumor cells **(Figure 6B)**. Such a set up would test the ability of biNTe to enhance the anti-myeloma activity of the CAR and limit immune escape even when half of the tumor cells lack expression of the CAR target. These myeloma-bearing mice were left untreated or were treated with 10^7^ FCRL5 CAR-iNKT or FcRL5 CAR iNKT plus BCMA biNTe. The combination of FCRL5 CAR-iNKT plus BCMA biNTe significantly prolonged disease progression to a greater extent than CAR-iNKT **(Figure 6C&D)**. Assessment of the fraction of FCRL5- and FCRL5+ myeloma cells in bone marrow and spleen at end-point confirmed the expected ratio of ∼1:1 in untreated animals while in FCRL5 CAR-iNKT plus BCMA biNTe and CAR-iNKT-treated animals this ratio was significantly higher at 10.1:1 and 5.1:1 respectively in favour of FCRL5-cells **(Figure 6E&F and Suppl Fig. 6A)**, consistent with a strong selective effect of FCRL5 CAR-iNKT against FCRL5+ disease. The overall disease burden in bone marrow and spleen was also significantly lower in CAR-iNKT plus BCMA biNTe compared to CAR-iNKT treated animals **(Figure 6G and Suppl Fig. 6B)** consistent with ability of biNTe to enhance the anti-myeloma activity of the CAR even when half of the tumor cells lack expression of the CAR target. In line with this, overall survival of FCRL5 CAR-iNKT plus BCMA biNTe-treated animals was significantly longer than that of animals treated with CAR-iNKT **(Figure 6H)**.

**Figure 6.**
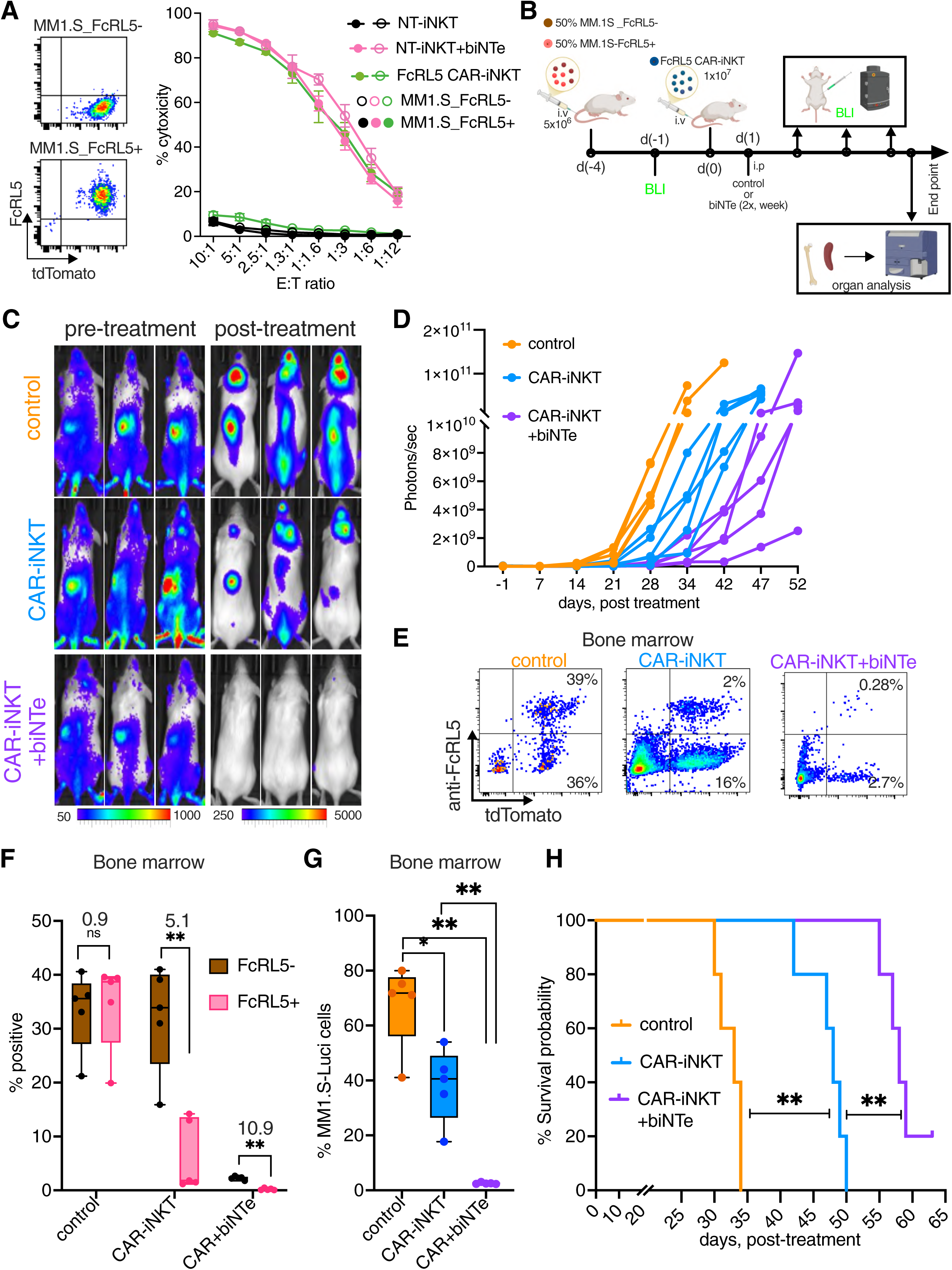
Enhanced anti-myeloma activity by dual CAR-iNKT plus biNTe targeting **A.** Flow cytometry plots show exogenous expression of FcRL5 in MM1.S (MM1.S_FcRL5+) cells and corresponding WT (MM1.S_FcRL5-) cells (left) and cytotoxic activity of FcRL5 CAR-iNKT or NT INKT cells incubated with or without biNTe against MM1.S_FcRL5- and MM1.S-FcRL5+ cells (right). (n=3). **B.** Schematic of *in vivo* experimental design associated with figure 6. **C&D.** Representative BLI images of mice from control and FcRL5 CAR-iNKT with or without biNTe group before and after treatment (week 4) (C) and their cumulative BLI data (D). (n=4-5 mice per group). **E&F.** Flow cytometry analysis at end point of the frequency of bone marrow MM1.S-FcRL5- and MM1.S-FcRL5+ cells in treated and control mice shown as dot plot (E) and cumulative data for each group (F). Numbers indicate the ratio between FcRL5- and FcRL5+ cells (n=4-5 mice per group). **G.** Flow cytometry analysis at end point of total MM1.S myeloma burden as determined by tdTomato fluorescence marker in the bone marrow of treated and control mice. **H.** Kaplan-Meier survival curve of myeloma-bearing mice left untreated (control) or treated with FcRL5 CAR iNKT cells or FcRL5 CAR-iNKT plus biNTe (n=4-5 mice per group) as shown in Figure 6B. ns-not significant, * p<0.05, ** p<0.01.

Together, these data provide strong proof-of-principle evidence that the combined CAR-iNKT plus biNTe modality immunotherapy enhances the therapeutic potency of the iNKT cell platform and can limit immune escape of tumor cells lacking the CAR target antigen in myeloma.

## Discussion

As iNKT cells emerge as a potentially more advantageous allogeneic immunotherapy platform than autologous or allogeneic T cells for cancer, there is a need to maximise their efficacy by optimising the therapeutic modules with which they are engineered. This would be particularly pertinent to multiple myeloma, an incurable blood cancer for which conventional autologous anti-BCMA CAR-T therapy may have potential to induce durable complete remissions in a small minority of patients^3^.

Here we describe a strategy to improve INKT-based ‘off-the-shelf’ cellular immunotherapy for MM first by optimising CAR design. CAR design, especially CD28z vs 4-1BBz endodomain choice, impacts efficacy and persistence of CAR-T cells by differential activation of downstream signalling pathways^19^. While CD28z supports robust T cell activation and more efficient targeting of CAR antigen low disease in part due to a lower level of trogocytosis by CAR-T cells of CAR antigen^34^ and robust phosphorylation of CD3z by LCK^35^, 4-1BBz promotes longer term persistence of CAR-T cells, in part through activation of non-canonical NFkB pathway^36^. Here, our systematic and multilayered comparison of five 2^nd^ and 3^rd^ generation CAR endodomain designs provides robust evidence that in the context of CAR-iNKT immunotherapy for myeloma a CD28z design would be the most advantageous in terms of potency without evidence of reduced persistence. Our data shows that a novel mechanism involving PLXND1-SEMA4A interaction determines differential avidity and thus higher anti- myeloma activity of CD28z CAR-iNKT as compared to CAR-iNKT with four different CAR endodomain designs. While engagement of PLXND1 on endothelial cells by SEMA4A has been shown to limit angiogenesis^27^, little is known on the role of PLXND1-SEMA4A in modulating function of immune cells. Plxnd1 through its interaction with Sema3E is required for migration of positively selected thymocytes into the medulla of the thymus^37^ while interaction of the Plxnd1-associated receptor Neuropilin 1 with Sema4A promotes transcriptional and functional stability of Tregs^38^. Of the potential PLXND1 ligands^25^, only SEMA4A is expressed in myeloma cells, hence its validation as a target for Ab- and CAR-based immunotherapy of multiple myeloma^28^. SEMA4A is also expressed in several other cancers and is required for checkpoint inhibitor-induced enhancement of anti-tumor T cell responses in lung cancer^39^, while Sema4A-expressing dendritic cells enhance activation and differentiation of antigen specific T cells via engagement of Tim2^40^. Additional work will be required to determine the signalling pathways that operate downstream of PLXND1 and underpin this advantage of CD28z CAR-iNKT.

Since both *FCRL5* and *SEMA4A* are chr1q genes and are overexpressed in myeloma cells with chr1q-amp, a critical adverse prognosis genetic feature, CD28z-based CAR-iNKT approach targeting FCRL5 in combination with BCMA could be particularly advantageous for this high risk myeloma^41^. This contrasts with the fact that both clinically licensed anti-BCMA CAR-T products for MM feature a 4-1BBz endodomain. However, since pre-clinical comparison of CD28z vs 4-1BBz FCRL5 CAR-T demonstrated the superior ability of the former to control myeloma *in vivo*^29^, whether 4-1BB would be the preferred endodomain for CAR-T treatment of MM remains an open question.

In terms of potency, consistent with previous work by us and others showing higher anti- tumor activity of CAR-iNKT in pre-clinical B cell lymphoma, high risk B ALL, and solid tumors^13–17^, we demonstrate that CAR-iNKT also outperform CAR-T in the context of multiple myeloma. This advantage of iNKT over T cells could be mediated both directly by the iTCR interacting with CD1d expressed on some tumors, including myeloma cells^12^, and engagement of activating NK receptors (e.g., NKG2D) by their ligands expressed by tumor cells; or indirectly- through enhancement of innate and adaptive anti-tumor immune responses, as well as targeting and depletion of CD1d-expressing immunosuppressive and tumor-promoting myeloid cells^10,15,16,42^. The latter mechanism, although not directly addressed in this work, would also be expected to contribute to the anti-tumor activity of CAR-iNKT cells in multiple myeloma given that tumor-associated myeloid cells have been shown to suppress effective anti-myeloma T cell responses and promote myeloma cell survival^43^.

Guided by pre-clinical and recent early clinical work demonstrating the anti-cancer potential of bispecific engagers administered in combination with adoptive transfer of effectors such as NK cells^44^ and to expand the options for multi-modular targeting of multiple myeloma we developed iNKT cell-specific engagers and demonstrated their anti-myeloma activity. Such engagers, with the exception of a γδTCR engager that cross-reacts with the TCRVα24Jα18 CDR3 idiotype of the iTCR^45^, have not been described. Our TCRVβ11-engaging bispecific antibody has two distinct properties: firstly, the option to target any tumor-associated antigen with one arm; and secondly, the availability of the TCRVα24Jα18 CDR3 loop to engage with CD1d-presented glycolipids thus further contributing to CD1d-dependent tumor killing. We envisage that iNKT engagers, with single or dual anti-tumor specificity, can be used flexibly in conjunction with other therapeutic modalities such as single or dual CAR and/or TCR thus achieving dual, triple or even quadruple targeting of myeloma, other blood or solid tumor cancers. Clinical development of this modular therapy would entail infusions of the iNKT cell engagers in conjunction with transfer of autologous or allogeneic, in vitro expanded iNKT or CAR/TCR-iNKT cells.

In summary, in the context of multiple myeloma, we provide proof-of-concept that iNKT engineered with optimally selected CAR design display superior anti-myeloma activity over CAR-T cells and their efficacy can be further broadened and enhanced by addition of iNKT cell engagers.

## Acknowledgments

KP was supported by Blood Cancer UK and the Kay Kendall Leukaemia Fund, KK was supported by an Imperial UKRI Impact Acceleration award, HR and BL were supported by Cancer Research UK, LR was supported by an MRC UKRI- AstraZeneca Fellowship. We also acknowledge support from Imperial NIHR Biomedical Research Centre.

## Contributions

KP established all CAR iNKT and performed *in vitro*, *in silico* and *in vivo* experiments and analysed data. KP & KK & YJ contributed on biNTe and FcRL5 CAR work. LR performed microscopy experiments and data analysis, KP, RN, IS & RR established and performed avidity assays. KP & AC performed RNA seq analysis. HR, IL, BL, MZ, VP, DL for technical support and MB, MA, AK provided patient material and KP, EB & ET performed SPR and data analysis and IR co-wrote the manuscript. AK and KP conceived the experimental plan and wrote the manuscript.

## Materials and Methods

### iNKT isolation and expansion, Cell culture, Primary cells

iNKT cells were isolated from healthy donor-derived peripheral blood mononuclear cells (PBMCs) sourced from NHS Blood and Transplant (NCI licence no. NCI2758) using anti-iNKT microbeads (cat no. 130-094-842; Miltenyi Biotec) followed by stimulation with irradiated PBMCs and anti-CD3 and anti-CD28 antibodies (cat no. 130-091-441; Miltenyi Biotec) and cultured in R10 media (RPMI 1640 supplemented with 10% fetal bovine serum and 1% Pen/strep, 4mM L-Glutamine, 1% non-essential amino acids, 1mM sodium pyruvate, 15mM Hepes buffer, 0.05mM 2-Mercaptoethanol) supplemented with 150U IL-15 (cat. no 130-095- 764; Miltenyi Biotec) and expanded as described^13,17^. The myeloma cell lines MM1.S and H929, which were sourced from ATCC, were cultured in RPMI 1640 supplemented with 10% fetal bovine serum and 1% Pen/strep, 1% L-Glutamine, 1% non-essential amino acids, 1% sodium pyruvate. Primary myeloma cells were collected under ethical committee approval (REC no. 11/H0308/9) from bone marrow samples derived from myeloma patients. The myeloma cells were idented by staining the cells with anti-human CD138 (Syndecan-1) (clone: MI15; BioLegend UK) and anti-human CD38 (clone: HIT2; BioLegend UK).

### Cloning, lentiviral transduction

BCMA CAR sequences were modified from Carpenter et al^23^, and modified by inserting (G4S)3 linker between light and heavy chain of scFV and appropriate costimulatory molecules and CD8a hinge and transmembrane and 7AA of cytoplasmic domain as shown in Figure 1A, and cloned into second generation plasmid containing SFFV promoter and WPRE. CAR lentiviral particles were packaged in 293T cells by transfection of 4.1µg CAR plasmid with packaging plasmids 5.4 µg psPAX2 and 2.9 µg pMD2.G plasmid for 48hrs. The viral particles were concentrated by ultracentrifugation at 23000 RPM, 4°C. The cells were transduced on 8µg/ml retronectin-coated plates by inoculating viral particles and cells at 1000g at 37°C.

#### CD28 BCMA CAR AA sequences

MALPVTALLLPLALLLHAARPDIVLTQSPPSLAMSLGKRATISCRASESVTILGSHLIHWYQQKPGQPPTLLIQLASNVQTGVPARFSGSGSRTDFTLTIDPVEEDDVAVYYCLQSRTIPRTFGGGTKLEIKGGGGSGGGGSGGGGSQIQLVQSGPELKKPGETVKISCKASGYTFTDYSINWVKRAPGKGLKWMGWINTETREPAYAYDFRGRFAFSLETSASTAYLQINNLKYEDTATYFCALDYSYAMDYWGQGTSVTVSSFVPVFLPAKPTTTPAPRPP TPAPTIASQPLSLRPEACRPAAGGAVHTRGLDFACDIYIWAPLAGTCGVLLLSLVITLYCNHRNRSKRSRLLHSDYMNMTPRRPGPTRKHYQPYAPPRDFAAYRSRVKFSRSADAPAYQQGQNQLYNELNLGRREEYDVLDKRRGRDPEMGGKPRRKNPQEGLYNELQKDKMAEAYSEIGMKGERRRGKGHDGLYQGLSTATKDTYDALHMQALPPR* (CD28 domain sequences)

#### 41BB domain

KRGRKKLLYIFKQPFMRPVQTTQEEDGCSCRFPEEEEGGCEL

#### OX40 domain

ALYLLRRDQRLPPDAHKPPGGGSFRTPIQEEQADAHSTLAKI

#### CD28-41BB domain

RSKRSRLLHSDYMNMTPRRPGPTRKHYQPYAPPRDFAAYRSKRGRKKLLYIFKQPFMRPVQTTQEEDGCSCRFPEEEEGGCEL

#### CD28-OX40 domain

RSKRSRLLHSDYMNMTPRRPGPTRKHYQPYAPPRDFAAYRSALYLLRRDQRLPPDAHKPPGGGSFRTPIQEEQADAHSTLAKI

These costimulatory domains were replaced in CD28 BCMA CAR to create all the BCMA CARs with different costimulatory molecules.

#### FcRL5 CAR AA sequences

MALPVTALLLPLALLLHAARPTGEVQLVESGPGLVKPSETLSLTCTVSGFSLTRFGVHWVRQPPGKGLEWLGVIWRGGSTDYNAAFVSRLTISKDNSKNQVSLKLSSVTAADTAVYYCSNHYYGSSDYALDNWGQGTLVTVSSGGGGSGGGGSGGGGSDIQMTQSPSSLSASVGDRVTITCKASQDVRNLVVWFQQKPGKAPKLLIYSGSYRYSGVPSRFSGSGSGTDFTLTISSLQPEDFATYYCQQHYSPPYTFGQGTKVEIKFVPVFLPAKPTTTPAPRPPTPAPTIASQPLSLRPEACRPAAGGAVHTRGLDFACDIYIWAPLAGTCGVLLLSLVITLYCNHRNRSKRSRLLHSDYMNMTPRRPGPTRKHYQPYAPPRDFAAYRSRVKFSRSADAPAYQQGQNQLYNELNLGRREEYDVLDKRRGRDPEMGGKPRRKNPQEGLYNELQKDKMAEAYSEIGMKGERRRGKGHDGLYQGLSTATKDTYDALHMQALPPR* (CD28 domain sequences)

### Flow cytometry and cytotoxicity assay

iNKT cells were stained with anti-human CD3 (clone UCHT1; BioLegend UK), anti-human TCR Valpha24-Jalpha18 (iNKT cell) (clone 6B11; BioLegend UK), anti-human TCR Vbeta11 (clone REA559; Miltenyi Biotec) anti-human CD45 (clone HI30; BioLegend UK), anti-human CD8 (clone SK1; BioLegend UK) and anti-human CD4 (clone:OKT4; BioLegend UK) for 30min at 4°C in FACS buffer (1x PBS+0.5% HSA). The cells were stained with 7AAD after washing and analysed using flow cytometry. PlexinD1 (cat no. FAB4160P; R&D systems), anti-human CD307e (FcRL5) (clone: REA391), anti-human BCMA (clone: REA315) antibodies were used.

CAR functional tests were performed by co-culturing CAR iNKT cells with target cells expressing CAR antigen at different E:T ratio in R10 media. The target cells were stained with cell trace violet (cat. no C34557; ThermoFisher), and the effector cells were depleted of cytokines overnight. After the indicated time of co-culture, the cells were washed with FACS buffer and cell viability was determined by staining the cells with 7AAD and flow cytometry. For primary myeloma cells, the total target cells were stained with cell trace violet before the coculture with NT iNKT cells, and the cells were stained with anti-CD138 antibodies to determine frequency of myeloma plasma cells after 20hours of culture and analysed using flow cytometry. The data were normalized to target cells alone control. The MM1.S cells were incubated with recombinant BCMA protein (clone: 10620-H40H-B-SIB) to block BCMA in biNTe experiment followed by incubation with iNKT cells (1:1) and biNTes at EC50 dose.

### Intracellular cytokine measurment

Non-transduced (NT) iNKT and BCMA CAR iNKT cells were co-cultured with MM1.S cells in the presence of 1x Brefeldin A overnight in R10 media without any cytokine stimulation. The cells were fixed and permeabilized by following standard manufacturer’s instructions (cat no. 420801 and 421002; BioLegend UK). The cells were stained with granzyme B (clone: GB11) and IFNγ (clone: 4S.B3; BioLegend UK) in FACS buffer for 30min, 4°C and measured in flow cytometry after washing twice with FACS buffer.

### Avidity assay

The iNKT cell avidity was determined using z-Movi ® Cell Avidity Analyzer (Lumicks/Lumicks CA BV). Briefly, 10^6^ cells/ml BCMA-expressing MM1.S or H929 cell monolayer was prepared on poly-L-lysine-coated temperature-controlled microfluidic chips (Lumicks/Lumicks CA BV) for 1hr. Simultaneously NT or CAR-modified iNKT cells were stained with cell trace far red dye (cat no. C34564; ThermoFisher) by following manufacturer’s instructions and incubated on the monolayer for 5 minutes. The indicated acoustic force was applied, and the fraction of bound cells was measured. For shear stress models, cell avidity was measured by applying shear-stress where MM1.S cell monolayer was prepared on 1µg/ml retronectin-coated ibid- treated slides for 1hr at 37°C. Simultaneously the effector iNKT cells were stained with cell trace green (cat no. C2925; ThermoFisher) or far-red dyes. The iNKT cells were incubated on the monolayer for 5 minutes and the indicated shear stress was applied continuously with an interval of 30 seconds between incremental shear stress levels. Total number of stained iNKT cells in the field was captured under fluorescence microscopy and the images were captured at every shear stress level. For PlexinD1 blocking, iNKT cells were incubated with either 1µg/ml isotype or anti-PLXND1 neutralizing antibody (cat no. PA5-47012; ThermoFisher) for 1 hr, 37°C and washed them in R10 media before applying them on the monolayer to measure the cell avidity efficiency as sated above. SEMA4A knock-out cells were generated by electroporating MM1.S cells with SEMA4A CRISPR gRNA and guide (156156422 CUCCAACAGGGGAUGAACGU – Modified) and SpCas9 (Synthego). The bound cells at all shear stress levels were counted using Image J software and normalized to bound cells at pre-shear stress application (0 dyne/cm^2^).

### Systemic and subcutaneous *in vivo* models of myeloma and bioluminescence

MM1.S cells were transduced with retrovirus to express luciferase gene and tdTomato, an enhanced, tandem-dimeric, monomeric red fluorescent protein derived from the original DsRed protein as a fluorescence marker (MM1.S-Luci). To establish a systemic myeloma model, 5-7×10^6^ MM1.S-luci cells were injected intravenously to engraft in 8-12 weeks old female NOD SCID Gamma (NSG) mice. The mice were randomized between the cages for different treatment. The mice were imaged for bioluminescent signal (BLI) to track the disease burden under 2% isoflurane (Zoetis UK)/medical oxygen administration via inhalation and one single dose injection of 150 mg/Kg D-Luciferin injection, potassium salt (cas no. 115144-35-9; Merk) in PBS via intraperitoneal (IP) route for 10 min before imaging the mice. The images were captured for up to 3 minutes at auto mode and medium binning of IVIS for both ventral and dorsal side of each mouse. The region of interest (ROI) data analysis was performed using Living Image software and expressed in radiance (unit of photons/s/cm^2^/sr) and the total BLI was calculated by adding both dorsal and ventral side signals. For subcutaneous model, 10^7^ MM1.S myeloma cells were injected with 200µl Matrigel® Basement Membrane Matrix (cat no. 354234; Corning) subcutaneously and the tumor size was measured using vernier calliper and tumor volumes were assessed using the formula (Volume = 1/2 * Length * Width²). For treatment, NT iNKT or CAR-iNKT cells or CAR-T cells were injected through tail vein intravenous route. For biNTe treatment, 5µg biNTe antibody per mouse was injected intraperitonially twice a week for up to 5 weeks. For PLXND1 blocking *in vivo*, the BCMA CAR iNKT treated group mice were injected with either 5µg Goat anti-human isotype or PLXND1 neutralizing antibody per mouse (cat no. PA5-47012; ThermoFisher) intraperitonially for 6 weeks, twice a week. The untreated and isotype control of BCMA treated group shown in Figure 3K were served as negative and positive control respectively shown in Figure 5L as these experiments were performed at the same period and were designed to reduce animal usage and to comply with 3R’s policy. Upon reaching humane end points of MM disease, the mice were sacrificed, and the organs were collected followed by cell preparation in FACS buffer and measured for tdTomato to track MM1.S cells in flow cytometry.

### RNA sequencing

iNKT cells were transduced with BCMA CARs and expanded as stated above after purification of BCMA CAR iNKT cells on day 5 post-transduction, to have >95% CARs for all the BCMA CARs, by staining the cells with recombinant BCMA-biotin (cat no. 10620-H40H-B-SIB; Stratech Scientific) followed by magnetically labelled anti-streptavidin PE beads-based purification by MACS magnetic column. The CAR and iNKT purity was assessed by flow cytometry before isolation of the total RNA, and RNA sequencing was performed after enriching for poly-A mRNA and sequenced by Novogene.

### Surface Plasmon Resonance (SPR)

Kinetics data were collected on Biacore S200 Biosensors (Cytiva). Soluble biotinylated BCMA were immobilised to Series S SA sensorchips, which were maintained with a continuous flow of PBS-P+ (20mM phosphate buffer with 2.7mM KCl, 137mM NaCl and 0.05% Surfactant P20). SA sensorchips were prepared as per Cytiva protocols; the sensor surface was conditioned with three consecutive one-minute injections of 1M NaCl in 50mM NaOH and washed with 70% isopropanol in 1M NaCl and 50mM NaOH after each ligand injection. The biotin tagged ligand was injected at 20ng/µL for 1800 seconds, reaching an RU of 3500. Kinetic data were collecting via SCK experiments with a 4-point dilution series (0.1, 1, 10, 200nM) in PBS-P+. The diluted compound series was injected in order of ascending concentration for 400s, across a specific flow cell. Dissociation was monitored during a 400s injection of assay buffer. An equivalent series of injections was carried out using Assay Buffer, which was used to correct flowcell-specific artefacts. Kinetics and affinity data were gathered using the Biacore Evaluation software.

### Bispecific antibody development

Monovalent biNTe composed of heterodimeric iTCR (clone: C21) scFV and anti-BCMA (clone: C11D5.3) scFV linked with Fc human wild type IgG1 with Fc-silent (L234A/L235A (LALA) and a ‘key-in-hole’ changes. A homodimeric IgG–scFv bispecific bivalent was constructed as the heavy chain (HC) encoded an anti-BCMA (C11D5.3) IgG (VH–CH1–hinge–CH2–CH3) C-terminally fused to an anti-Vβ11 scFv (C21) via a (G₄S)₃ linker. The light chain (LC) encoded the matched anti-BCMA (C11D5.3) VK–CK. HC and LC were cloned in separate mammalian expression vectors. The biNTes were expressed by transiently co-transfecting 30µg of both plasmids of dimeric biNTe with 180µg PEI (cat no. 7854; biotechne) (PEI:DNA 3:1 w/w mixed in 2ml plain DMEM media) in 293T cells at 85% confluence grown in T175 flask. The media was replaced with 30ml fresh media after 24hours and added additional 30ml fresh media replaced with 30 mL fresh complete DMEM (10% FBS, penicillin/streptomycin, glutamine) after 24 hours and added additional 30 mL complete medium after 48hours. Cultures were maintained at 37 °C, 5% CO₂. On Day 6, culture supernatants were collected and clarified by two sequential centrifugations. The clarified supernatant was then filtered through a 0.4 µm membrane The engagers were purified using rProtein A/Protein G GraviTrap column (cat no. GE28-9852-56; Cytiva) by passing 60ml of culture supernatant followed by washing twice with 10ml 1xPBS. The antibody was eluted in 1ml fractions of 0.1M Glycine-HCL (pH 2.7) elution buffer and neutralized by 60µl 1M Trizma hydrochloride solution pH 9.0 (cat no. T2819; Merk) and the elution was repeated for 6 times. The eluted biNTe were concentrated using Amicon Ultra-15 centrifugal filters (Diameter (Metric): 29.7 mm MWCO: 30,000 d) (cat no. UFC903024; Merk) and washed twice with 1x PBS. The purity of biNTe was assessed by SDS-PAGE under reducing and non-reducing conditions.

#### iTCR scFV AA sequences

DIKMTQSPSSMYASLGERVTITCKASQDINSYLSWFQQKAGKSPKTLIYRANRLVDGVPSRFSGSGSGQDYSLTISSLEYEDMGIYYCLQYDEFPFTFGGGTRLEIKGGGGSGGGGSGGGGSQVQLQQSGPEVVRPGVSVKISCKGSGYRFTDSAMHWVKQSHAKSLEWIGVISSYNGNTNYNQKFKGKATMTVDKSSSTAYMELARMTSEDSAIYYCARSRDAMDYWGQGTSVTVSS

#### anti-BCMA scFV AA sequences

DIVLTQSPPSLAMSLGKRATISCRASESVTILGSHLIHWYQQKPGQPPTLLIQLASNVQTGVPARFSGSGSRTDFTLTIDPVEEDDVAVYYCLQSRTIPRTFGGGTKLEIKGGGGSGGGGSGGGGSQIQLVQSGPELKKPG ETVKISCKASGYTFTDYSINWVKRAPGKGLKWMGWINTETREPAYAYDFRGRFAFSLETSASTAYLQINNLKYEDTATYFCALDYSYAMDYWGQGTSVTVSS

#### Monovalent (anti-BCMA scFV-IgG) AA sequences

MPLLLLLPLLWAGALADIVLTQSPPSLAMSLGKRATISCRASESVTILGSHLIHWYQQKPGQPPTLLIQLASNVQTGVPARFSGSGSRTDFTLTIDPVEEDDVAVYYCLQSRTIPRTFGGGTKLEIKGGGGSGGGGSGGGGSQIQLVQSGPELKKPGETVKISCKASGYTFTDYSINWVKRAPGKGLKWMGWINTETREPAYAYDFRGRFAFSLETSASTAYLQINNLKYEDTATYFCALDYSYAMDYWGQGTSVTVSSGGGGSGGGGSGGGGSDKTHTCPPCPAPEAAGGPSVFLFPPKPKDTLMISRTPEVTCVVVDVSHEDPEVKFNWYVDGVEVHNAKTKPREEQYNSTYRVVSVLTVLHQDWLNGKEYKCKVSNKALPAPIEKTISKAKGQPREPQVYTLPPCRDELTKNQVSLWCLVKGFYPSDIAVEWESNGQPENNYKTTPPVLDSDGSFFLYSKLTVDKSRWQQGNVFSCSVMHEALHNHYTQKSLSLSPG

#### Monovalent (iTCR scFV-IgG) AA sequences

MPLLLLLPLLWAGALADIKMTQSPSSMYASLGERVTITCKASQDINSYLSWFQQKAGKSPKTLIYRANRLVDGVPSRFSGSGSGQDYSLTISSLEYEDMGIYYCLQYDEFPFTFGGGTRLEIKGGGGSGGGGSGGGGSQVQLQQSGPEVVRPGVSVKISCKGSGYRFTDSAMHWVKQSHAKSLEWIGVISSYNGNTNYNQKFKGKATMTVDKSSSTAYMELARMTSEDSAIYYCARSRDAMDYWGQGTSVTVSSGGGGSGGGGSGGGGSDKTHTCPPCPAPEAAGGPSVFLFPPKPKDTLMISRTPEVTCVVVDVSHEDPEVKFNWYVDGVEVHNAKTKPREEQYNSTYRVVSVLTVLHQDWLNGKEYKCKVSNKALPAPIEKTISKAKGQPREPQVYTLPPCRDELTKNQVSLTCLVKGFYPSDIAVEWESNGQPENNYKTTPPVLDSDGSFFLTSKLTVDKSRWQQGNVFSCSVMHEALHNHYTQKSLSLSPG

#### Anti-BCMA FAB AA sequences

Heavy chain: QIQLVQSGPELKKPGETVKISCKASGYTFTDYSINWVKRAPGKGLKWMGWINTETREPAYAYDFRGRFAFSLETSASTAYLQINNLKYEDTATYFCALDYSYAMDYWGQGTSVTVSS Light chain: DIVLTQSPPSLAMSLGKRATISCRASESVTILGSHLIHWYQQKPGQPPTLLIQLASNVQTGVPARFSGSGSRTDFTLTIDPVEEDDVAVYYCLQSRTIPRTFGGGTKLEIK

#### Fc IG AA sequences

ASTKGPSVFPLAPSSKSTSGGTAALGCLVKDYFPEPVTVSWNSGALTSGVHTFPAVLQSSGLYSLSSVVTVPSSSLGTQTYICNVNHKPSNTKVDKKVEPKSCDKTHTCPPCPAPEAAGGPSVFLFPPKPKDTLMISRTPEVTCVVVDVSHEDPEVKFNWYVDGVEVHNAKTKPREEQYNSTYRVVSVLTVLHQDWLNGKEYKCKVSNK ALPAPIEKTISKAKGQPREPQVYTLPPSRDELTKNQVSLTCLVKGFYPSDIAVEWESNGQPENNYKTTPPVLDSDGSFFLYSKLTVDKSRWQQGNVFSCSVMHEALHNHYTQKSLSLS

### Immunofluorescence and confocal microscopy

For immune synapse imaging, 10^6^ /ml of untransduced iNKT or BCMA CAR-iNKT cells were mixed with MM1.S target cell at 1:1 ratio in serum-free RPMI and incubated for 30 to 45min at 37°C on a glass microscope slide (#PH299 BK, Hendley Essex). Cells were fixed in ice-cold methanol for 5min, blocked in 2% BSA in PBS for 40 minutes and stained overnight at 4°C with rabbit anti-actin (cat no. A2066, MilliporeSigma) and mouse anti-γTubulin (cat no. T6557, MilliporeSigma) primary antibodies. The next day, samples were washed and incubated with a donkey anti-rabbit (cat no. A-31572, ThermoFisher) or a goat anti-mouse (cat no. A-11001, ThermoFisher) fluorophore-conjugated secondary antibody. For PLXND1 expression analysis, iNKT cells only were incubated on the slides for 15min at 37°C, fixed for 15 min in 4% paraformaldehyde (cat no. 15710-S, Electron Microscopy Systems) and permeabilized in 0.1% Triton X-100 (Sigma) for 5min. Samples were then blocked and stained as described above using a primary anti-Plexin D1 antibody (cat no. ab28762; Abcam) followed by a goat anti-rabbit fluorophore-conjugated secondary antibody (cat no. A-21428; ThermoFisher) and Alexa Fluor® 488 phalloidin (cat no. 8878; Cell Signaling Technoloy) for 45min at RT. All samples were mounted in antifade medium with DAPI (cat no. H-1500-10; Vector Laboratories) and were acquired on an LSM-780 confocal laser scanning microscope. Z-stack series of 0.62 μm intervals were taken with a ×63 oil DIC Plan-Apochromat objective (numerical aperture 1.4).

### Data analysis and software

All the flow cytometry analysis was performed using Flow Jo software. The graphical data and statistics were performed using Prism 10 software and Biorender tools. Non-parametric Mann-Whitney t test used to compare independent groups and significance were calculated based on the p values. All the RNA sequencing traces were aligned using STAR and differential analysis performed using DESeq2 R Bioconductor packages by paired analysis. Image J was used for automatic cell counting of bound cells on monolayers for cell avidity experiments. In-vivo Imaging system (IVIS) and Aura software were used for *in vivo* bioluminescence photons analysis.

---

**Supplementary Figure 1 (related to Figure 1).**
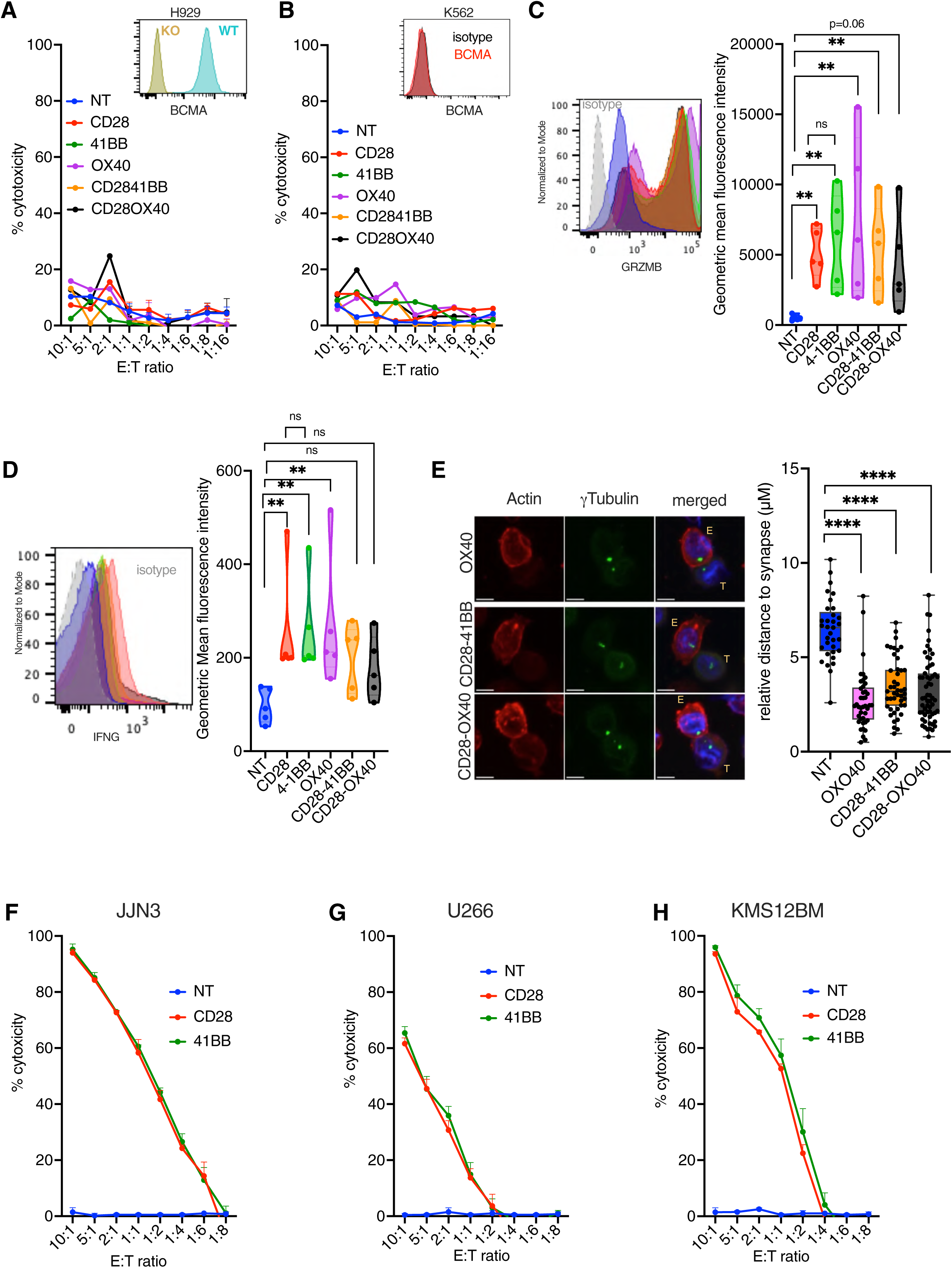
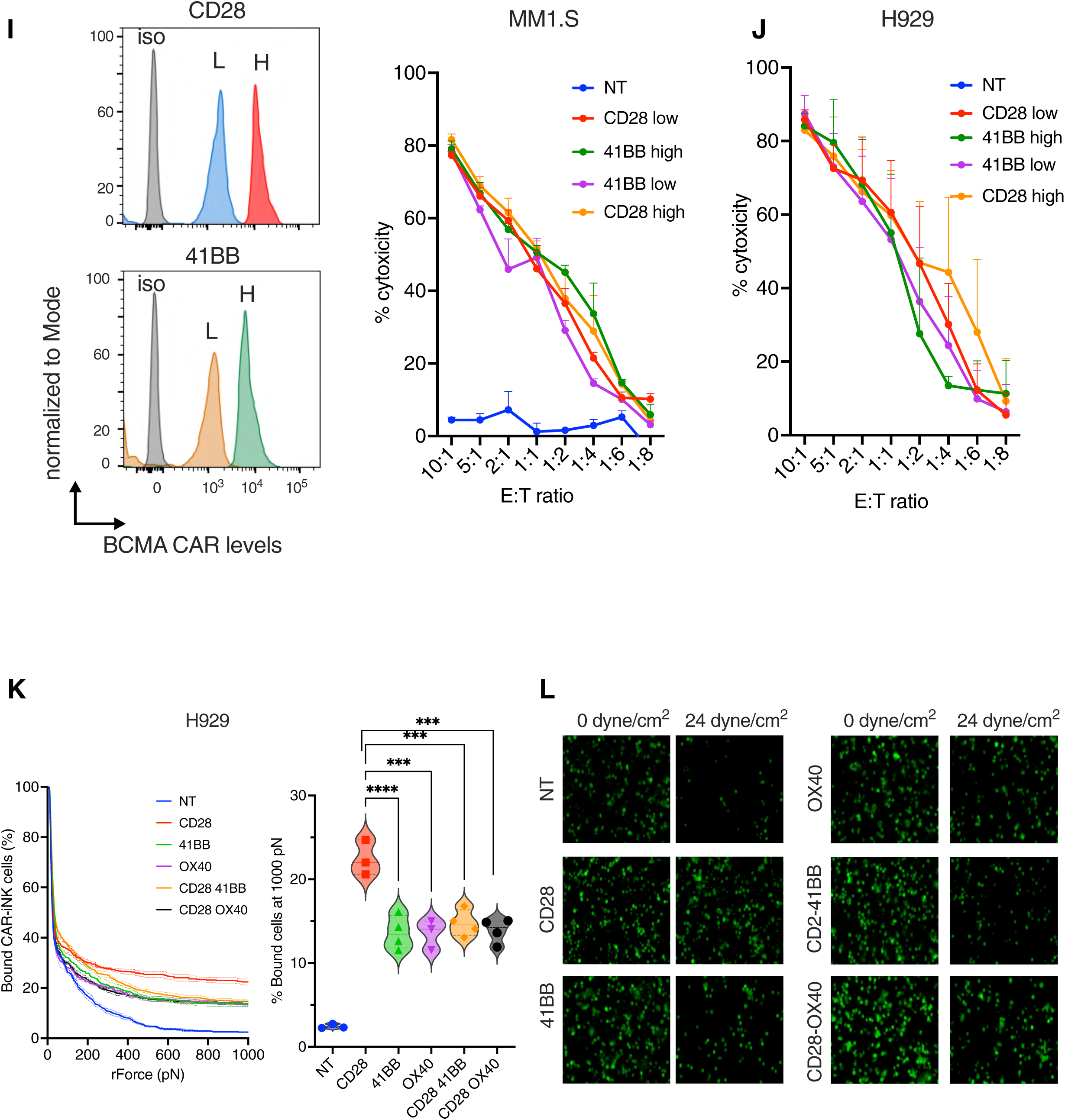
**A.** BCMA expression in BCMA H929 WT and BCMA KO cells shown in histogram and their cytotoxicity by CAR iNKT cells. **B.** BCMA expression in K562 erythromyeloid cells shown in histogram and CAR iNKT cell-mediated cytotoxicity against the same cells **C.** Left: Intracellular expression of granzyme B levels in CAR-iNKT measured by flow cytometry after exposing cells to the MM1.S myeloma cells for 16hours. Right: cumulative data from four different donors. **D.** Left: Intracellular expression of interferon gamma levels in CAR-iNKT measured by flow cytometry after exposing the cells to the MM1.S myeloma cells for 16hours. Right: cumulative data from four different donors. **E.** Immune synapse formation between the shown BCMA CAR iNKT cells and target MM1.S cells as assessed by confocal microscopy and measured by the distance of centrosome to the synapse as identified by anti-actin and -g-tubulin staining (data from three different donors; each point represents a synapse point distance). **F-H.** Comparison of cytotoxic activity of CD28z and 41BB CAR iNKT cells against JJN3 (F), U266 (G) and KMS12BM (H) myeloma cells is shown. **I-J.** After CD28z and 41BBz BCMA CAR iNKT cells were FACS-sorted for low (L) and high (H) levels of CAR expression (I); their cytotoxic activity against MM1.S cells (I) and H929 cells (J) was tested. iso-isotype, L-low BCMA CAR iNKT, H-high BCMA CAR iNKT, NT-non-transduced iNKT cells. **K.** Left: CD28 CAR iNKT cell binding activity on H929 myeloma cells under increasing levels of acoustic force as compared to all other CAR- and NT iNKT; right: CAR-iNKT binding at rForce 1000 pN (data representative of n=2 donors). **L.** Representative images of cell binding before shear stress (0 dyne/cm^2^) and after 24 dyne/cm^2^ which are related to Figure 1I. ns-not significant, ** p<0.01, **** p< 0.0001.

**Supplementary Figure 2 (related Figure 2).**
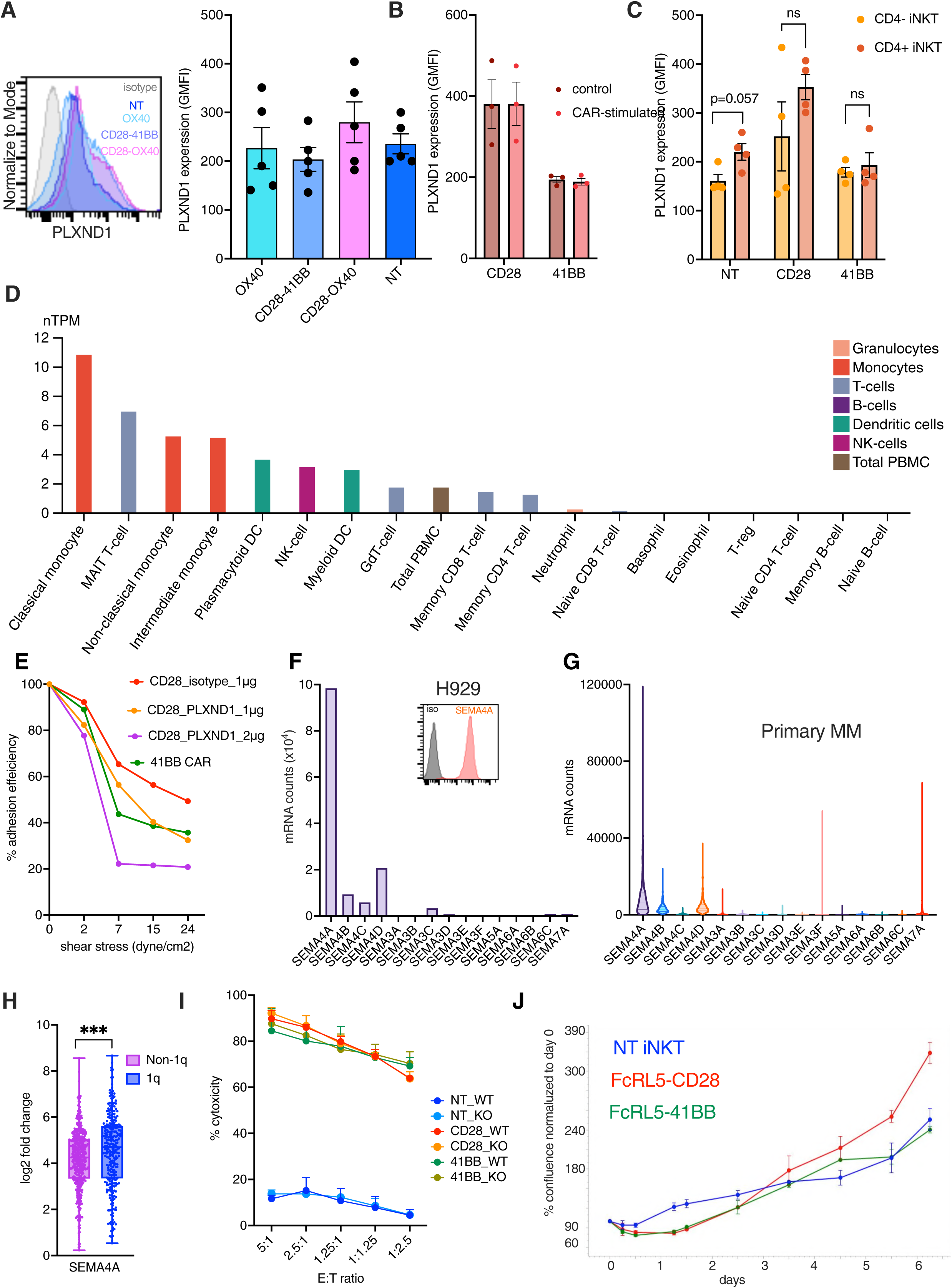
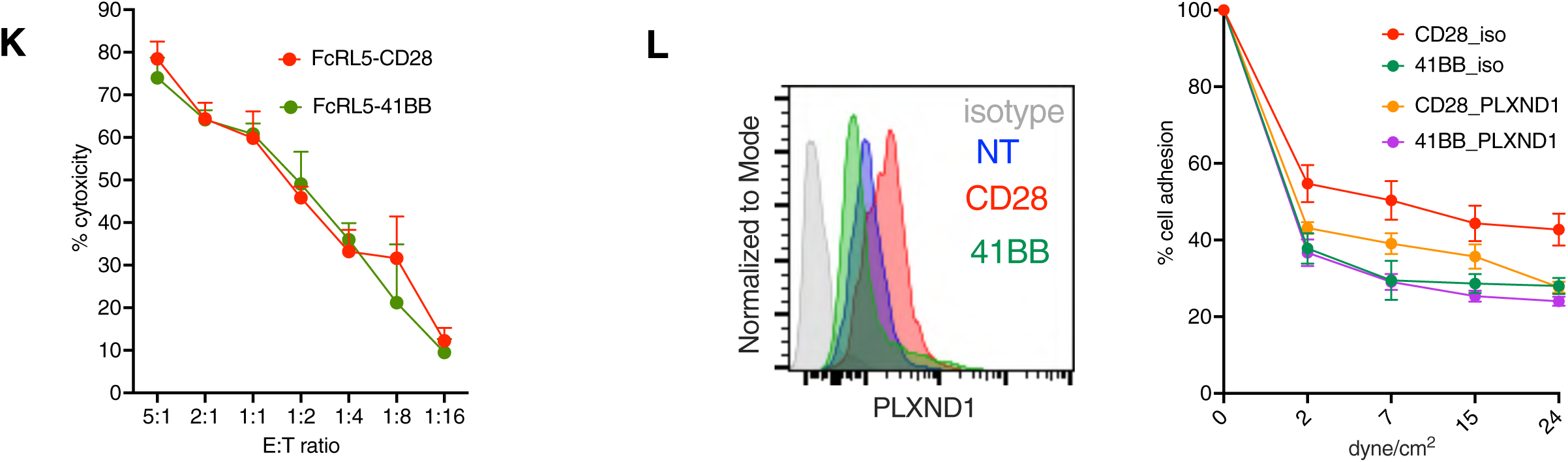
**A.** Expression of PLXND1 in shown CAR iNKT cells with different co-stimulatory molecules. Left histogram: flow-cytometric assessment; right: cumulative data from n=5 donors. **B.** PLXND1 expression in CD28 CAR iNKT cells as assessed by flow-cytometry after stimulation with MM1.S myeloma cells for 24hr (n=3). **C.** PLXND1 expression in CD4+ and CD4- fractions of iNKT cells as assessed by flow-cytometry ad GMFI (n=4). **D.** Expression of PLXND1 in subpopulations of immune cells. Data derived from Human Protein Atlas database. **E.** Dose-dependent reduction of avidity of BCMA CAR iNKT with CD28z co-stimulatory molecules by PLXND1 blocking antibody. BCMA CAR with 4-1BBz shown as control. **F.** SEMA4A mRNA count values and its family members in H929 cells (CLLE Dataset). The histogram shows SEMA4A expression on the surface of H929 cells measured by flow cytometry. **G.** Expression of SEMA4A mRNA counts and its family members in 770 primary myeloma cells from the MMRF CoMMpass study. **H.** SEMA4A expression is rhigher in Chr 1q+ primary MM cells as compared to non 1q MM cells (MMRF CoMMpass Dataset). **I.** No difference in cytotoxicity by BCMA CAR iNKT cells observed against SEMA4+ MM1.S (WT) and SEMA4- MM1.S (KO) cells (n=3 donors). **J.** FcRL5-CD28z CAR iNKT cells shows higher proliferative potential than its 41BBz counterpart and NT iNKT cells in short-term proliferation assay in the presence of IL-15 as assessed by real- time IncuCyte®imaging (n=3 donors). **K.** No difference in cytotoxic activity between FcRL5-CD28z and FcRL5-41BBz CAR iNKT cells against MM1.S-FcRL5+ target cells. **L.** Left: flow cytometry analysis of Plexin D1 expression in FcRL5 CAR iNKT cells with CD28z or 41BBz costimulatory molecules as compared to NT iNKT cells. Right: cell avidity of FcRL5- CD28z and FcRL5-4-1BBz CAR iNKT cells on MM1.S-FcRL5+ cells under shear stress with isotype control or PLXND1-blocking antibody at 1µg/ml concentration (n=3). ns-not significant, *** p<0.001.

**Supplementary Figure 3 (related Figure 3).**
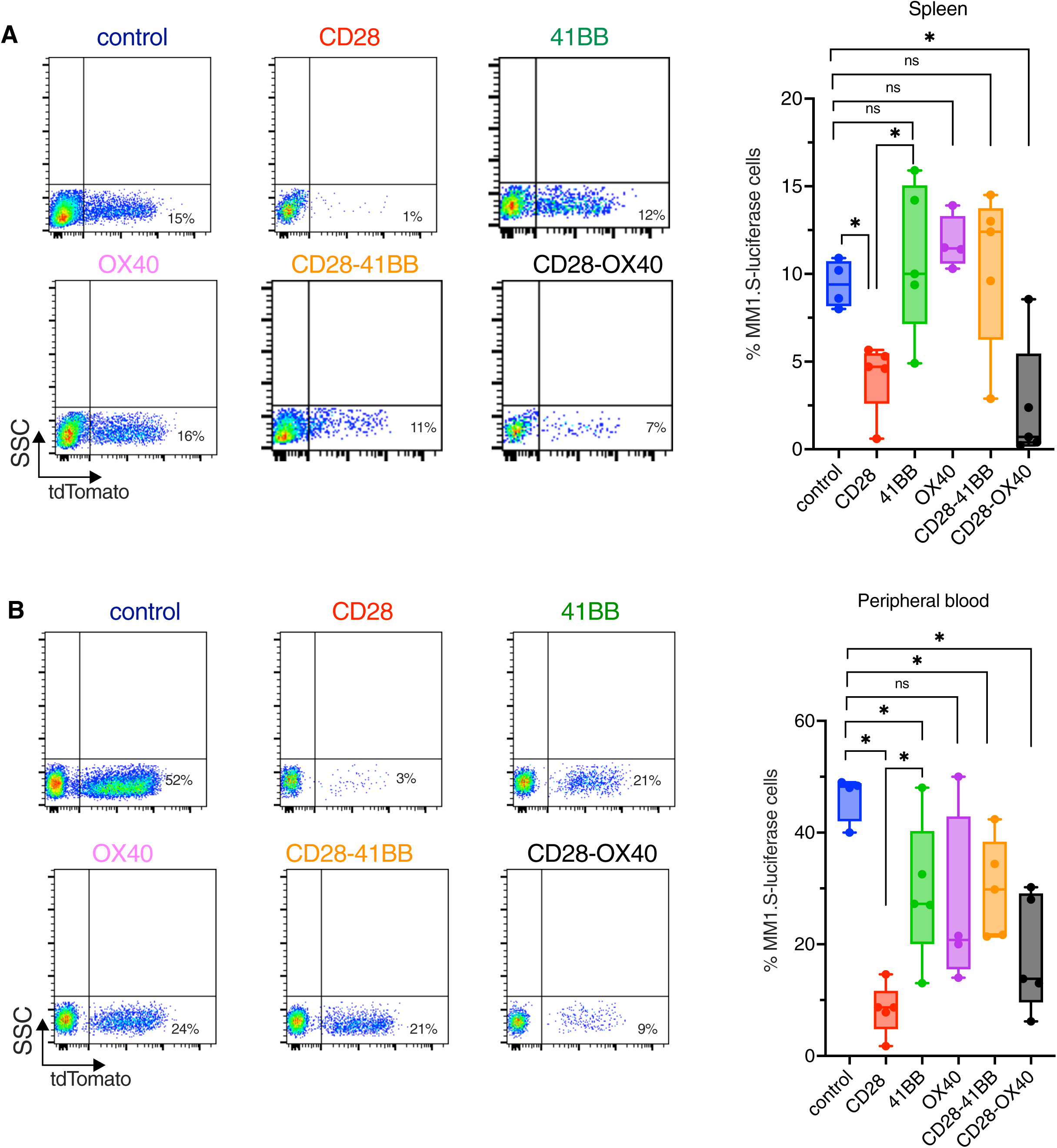
**A-D.** Frequency of MM1.S-luci tumor cells in the spleen (A), peripheral blood at end-point (n=4-5 mice per group) as measured by tdTomato fluorescence marker using flow cytometry. ns-not significant, * p<0.05, ** p<0.01.

**Supplementary Figure 4 (related for Figure 4).**
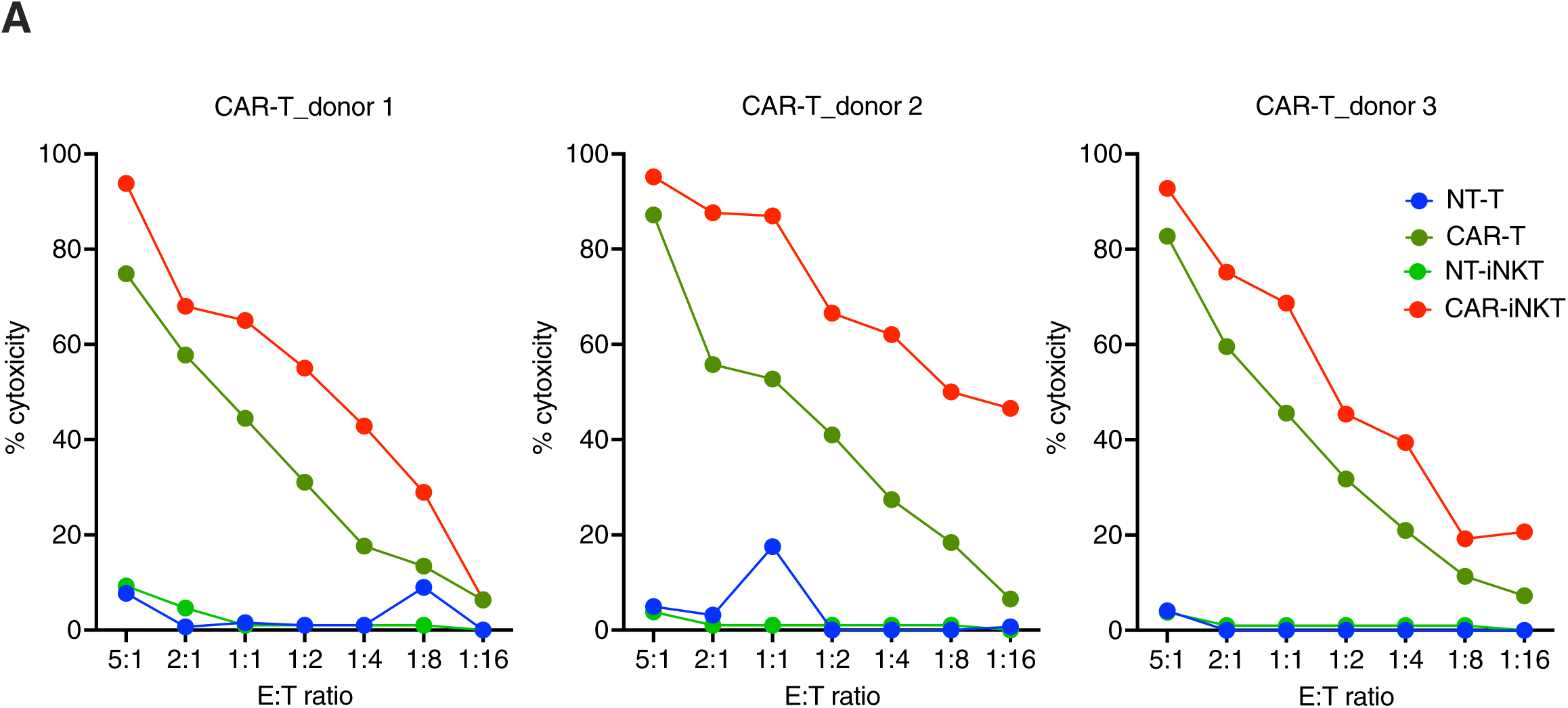
**A.** Cytotoxic potential of same donor BCMA-CD28z CAR iNKT and BCMA-CD28z CAR-T from 3 different donors against MM1.S cells.

**Supplementary Figure 5 (related to Figure 5).**
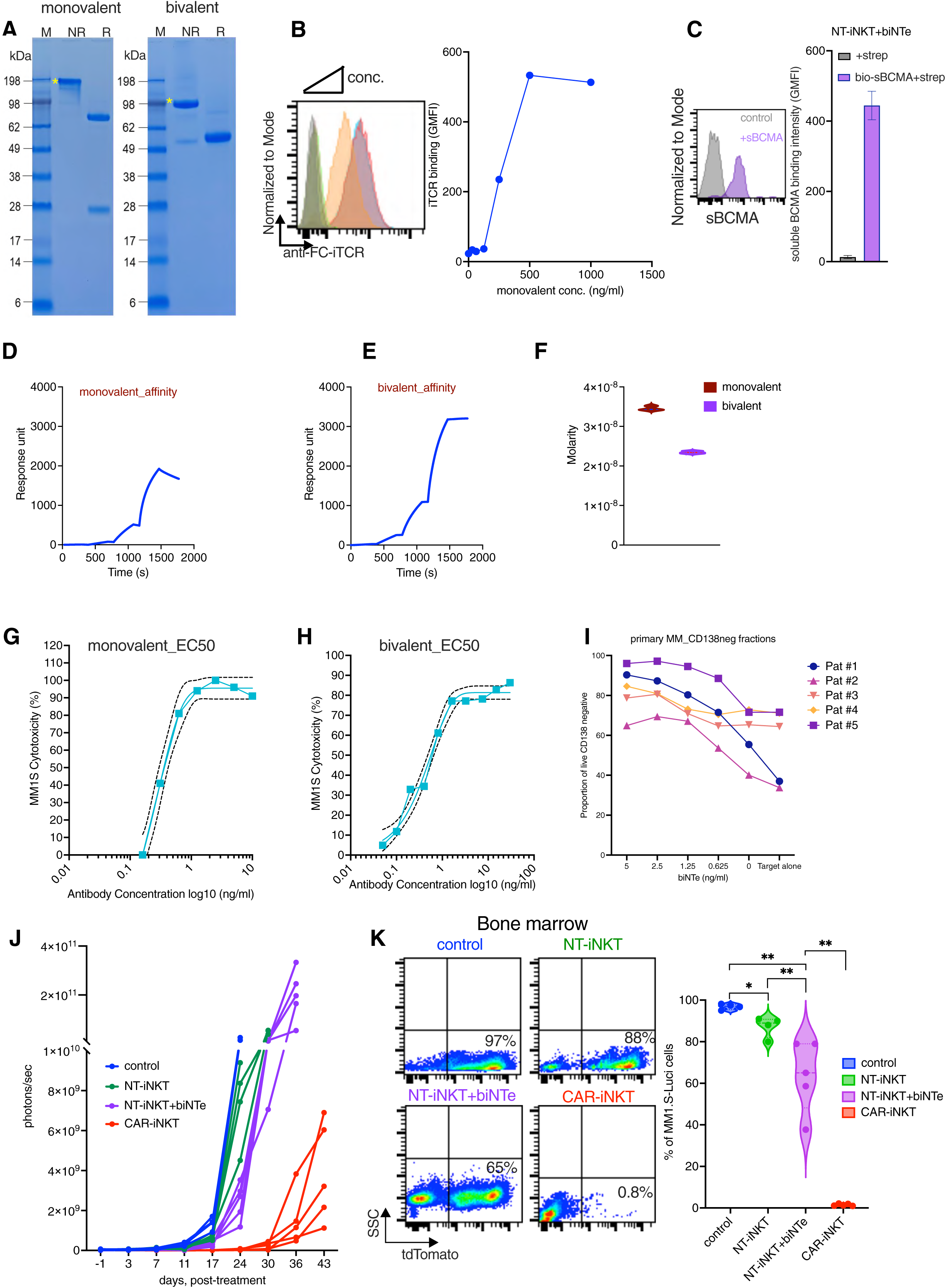
**A.** SDS-PAGE gel analysis of purity of monovalent (left) and bivalent biNTe (right) under reducing (R) and non-reducing (NR). M indicates marker. * Indicates the band size ∼198.9 kDa for monovalent and ∼78.9 kDa for bivalent biNTe under non-reducing conditions. **B.** Increasing levels of binding efficiency at increasing level of antibody concentration (0-1µg) for monovalent antibody to iNKT cells. Concentration range (0-1µg/ml). **C.** Flow-cytometric analysis of pure iNKT cells after staining with bivalent biNTe and fluorescently labelled streptavidin (control: grey) or with bivalent biNTe, biotinylated soluble BCMA and fluorescently labelled streptavidin (purple). Representative histogram (left) and its cumulative data (right) shown as GMFI. (n=3 donors). **D-F.** Affinity strength of monovalent (D), bivalent (E) antibodies on soluble BCMA against indicated time and their kd values (F) as determined in SPR assay, (n=3). **G&H.** EC50 values shown for monovalent (G) and bivalent engagers (H) against MM1.S by co-culture of NT iNKT at 1:1 E:T ratio plus indicated engagers. EC50 values were calculated by using non-linear fit, log(agonist) vs. response function in prism 10 software. **I.** CD138 negative fractions of primary myeloma cells after incubating with NT iNKT cells and biNTe as part of the CD138+ fractions analysis shown in Figure 5G. **J.** Cumulative BLI data for mice treated with NT iNKT and/or biNTe, and CAR iNKT and control group as shown in Figure 5J. (n=4-5 mice per group). **K.** Flow-cytometric analysis for frequency of MM1.S cells in the bone marrow from each group at end point. Left: representative flow-cytometry plots; right: cumulative data. (n=4-5 mice per group). * p<0.05, ** p<0.01

**Supplementary Figure 6 (related to Figure 6).**
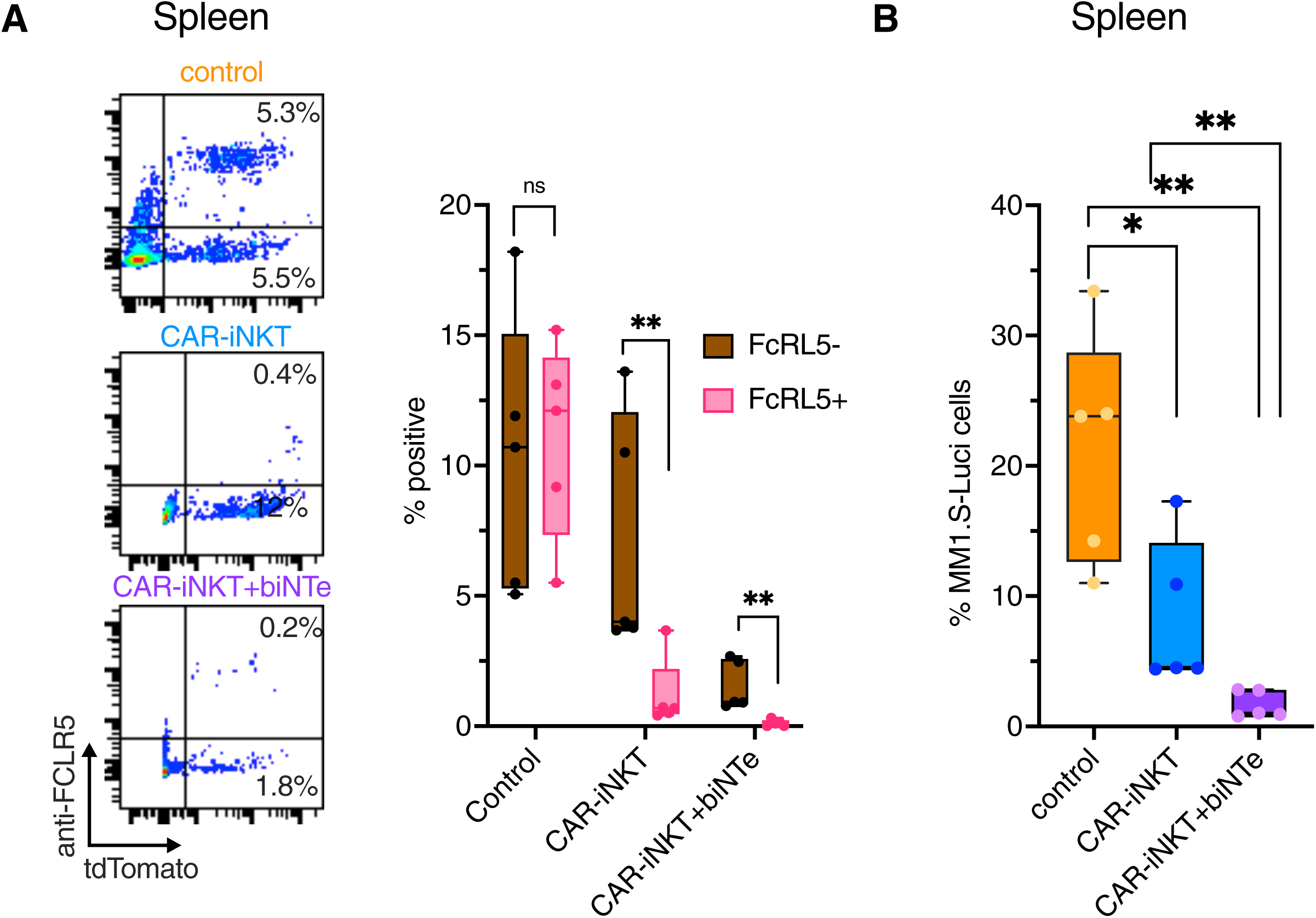
**A.** Flow cytometry analysis of frequency of MM1.S_FcRL5- and MM1.S-FcRL5+ cells in the spleen of treated and control mice at the end point shown as dot plot (left) and cumulative data for each group (right). (n=4-5 mice per group). **B.** Flow cytometry analysis of total MM1.S myeloma burden as determined by tdTomato fluorescence marker in the spleen of treated and control mice at the end point. (n=4-5 mice per group). **C.** Left: Flow cytometry analysis for frequency of MM1.S_FcRL5- and MM1.S_FcRL5+ cells proportion in the extramedullary tumor of FCRL5 CAR-iNKT plus biNTe treated group (n=5 mice). ns-not significant, * p<0.05, ** p<0.01.

