## Supplementary material for "Enhancing the iNKT cell immunotherapy platform by combining optimised CAR endodomains with novel iNKT engagers": Suppl Table 1

Common genes differentially expressed between groups

| NT vs CD28 | log fold change | padj |
|---|---|---|
| SLC22A17 | 6.437448317 | 0.03458404 |
| SPINK2 | 5.483968252 | 0.00051695 |
| ITGAD | 4.113349959 | 1.14E-05 |
| LINC01229 | 4.031022431 | 0.00015232 |
| ADCY6 | 3.648054936 | 3.91E-10 |
| ADAMTS10 | 3.19867216 | 0.00430368 |
| DAPK1 | 3.151737941 | 0.00085486 |
| HRH2 | 3.146399899 | 0.00430368 |
| PITPNM2 | 2.9184052 | 1.79E-05 |
| BTLA | 2.617527811 | 0.00388383 |
| SAMD14 | 2.499666405 | 0.02002882 |
| FOXD1 | 2.487780058 | 0.04460047 |
| AXIN2 | 2.452950797 | 0.04859656 |
| PCGF2 | 2.358241077 | 0.02368751 |
| KRT86 | 2.344622798 | 0.02002882 |
| ME3 | 2.330107667 | 0.02002882 |
| PTGIS | 2.325981492 | 0.02002882 |
| KRT81 | 2.224798334 | 0.0046832 |
| BAIAP3 | 2.219063573 | 0.00455536 |
| CDCP1 | 2.063294074 | 0.04325102 |
| RUSC2 | 2.025786184 | 0.04438752 |
| PLPP1 | 1.994218142 | 0.00135404 |
| PDE7B | 1.993811859 | 0.0324587 |
| GSTA4 | 1.933655607 | 0.00388383 |
| SPNS2 | 1.898788236 | 0.0072805 |
| SIRPG | 1.749994302 | 0.00135404 |
| NMUR1 | 1.727607887 | 0.00181187 |
| SOGA1 | 1.669905136 | 0.00388383 |
| CDH24 | 1.635608416 | 0.02612201 |
| PALLD | 1.54638735 | 0.0090473 |

| NT vs 41BB | log fold change | padj |
|---|---|---|
| LURAP1L | 6.791970948 | 0.01328462 |
| SLC22A17 | 6.719726872 | 0.01628337 |
| EPB41L4B | 6.337989459 | 0.02339219 |
| ASAP2 | 6.12813308 | 0.02747521 |
| TIE1 | 6.112332162 | 0.00099041 |
| COL6A2 | 6.056316525 | 2.25E-07 |
| MOXD1 | 5.774803881 | 7.08E-05 |
| MMP9 | 5.672551312 | 1.84E-11 |
| MB | 5.621199339 | 0.00426851 |
| COL6A1 | 5.441063747 | 1.84E-11 |
| PTPN3 | 3.723313092 | 0.04690818 |
| HMSD | 3.690105584 | 0.01147644 |
| LHFPL6 | 3.644160025 | 0.01328462 |
| PAWR | 3.625229132 | 0.02699351 |
| KRT87P | 3.611866188 | 0.0036069 |
| ADCY6 | 3.447844915 | 2.33E-09 |
| NLGN4Y | 3.403382651 | 0.01127803 |
| EBI3 | 3.397184001 | 0.00039331 |
| TLCD5 | 3.3773161 | 0.04343992 |
| CTTNBP2NL | 3.052324101 | 0.00827437 |
| ITGAD | 2.925590666 | 0.01029495 |
| SOX4 | 2.924739723 | 0.02186817 |
| KRT7 | 2.903579711 | 5.64E-10 |
| SOX18 | 2.68779768 | 0.00973163 |
| KRT86 | 2.545734885 | 0.00396641 |
| PTGS2 | 2.531896635 | 0.00023744 |
| RUSC2 | 2.50459385 | 0.00189913 |
| NTRK1 | 2.475962184 | 3.70E-06 |
| P2RY14 | 2.438463521 | 0.03248455 |
| PLA2G4A | 2.402182458 | 0.00124586 |

| | | |
|---|---|---|
| KRT7 | 1.542808659 | 0.03330275 |
| H1-0 | 1.452111307 | 0.03843425 |
| BACE2 | 1.384142029 | 0.02763009 |
| SERPINB6 | 1.295263327 | 0.02215584 |
| ASPH | 1.293238408 | 0.02215584 |
| CLIP2 | 1.200355271 | 0.0233335 |
| TPCN1 | 1.12132177 | 0.02002882 |
| HDAC5 | 1.087867494 | 0.0438578 |
| FCMR | 1.087289616 | 0.00062306 |
| MCOLN2 | 1.014813604 | 0.02002882 |
| HSPB1 | 0.957387898 | 0.02612201 |
| LPGAT1 | -0.795601695 | 0.00578245 |
| NETO2 | -0.891697225 | 0.02002882 |
| IFNG | -1.659564538 | 0.00058421 |
| RHOU | -1.909714566 | 0.0233335 |
| GZMH | -2.079510026 | 0.03013052 |
| LUCAT1 | -2.253723137 | 0.01997764 |
| PLXDC2 | -2.508820634 | 0.03013052 |
| SLC2A6 | -2.630583493 | 9.66E-07 |

| | | |
|---|---|---|
| SAMD14 | 2.399797138 | 0.02392053 |
| DAPK1 | 2.396298512 | 0.03808316 |
| NR5A2 | 2.345738369 | 0.01849229 |
| NBL1 | 2.276814131 | 8.04E-10 |
| RTN4RL1 | 2.274342501 | 0.04690818 |
| GPR35 | 2.25786775 | 3.31E-06 |
| PCGF2 | 2.23921973 | 0.03435364 |
| KRT81 | 2.211851944 | 0.00386403 |
| PALLD | 2.201296849 | 7.46E-07 |
| ICOSLG | 2.1864134 | 0.01628337 |
| EVC | 2.158659033 | 2.33E-09 |
| MSC | 2.12920707 | 0.04946625 |
| MSC-AS1 | 1.971026618 | 0.03556578 |
| FBN1 | 1.963584251 | 0.01029495 |
| PANX2 | 1.813879409 | 0.00232295 |
| EPB41L2 | 1.724151719 | 0.02554069 |
| GATA3-AS1 | 1.623218123 | 0.01849229 |
| C20orf204 | 1.60134153 | 0.00386403 |
| BIRC3 | 1.594366018 | 2.71E-05 |
| ASB2 | 1.216209171 | 0.00237567 |
| DUSP10 | 1.171524825 | 0.00124586 |
| DLG4 | 1.052859488 | 0.03435364 |
| HSPB1 | 1.020037102 | 0.00909481 |
| HIC1 | 0.975776137 | 0.04946625 |
| DNPH1 | 0.915003531 | 0.00630772 |
| CHCHD10 | 0.888049635 | 0.02094162 |
| SERTAD2 | 0.853813613 | 0.03958587 |
| TC2N | -0.75378948 | 0.04009745 |
| TRIB2 | -0.863061506 | 0.02612283 |
| NPC2 | -0.885536335 | 0.00630772 |
| SAMD3 | -1.118263909 | 0.01423645 |
| HS3ST3B1 | -1.285949514 | 0.01808374 |

| | | |
|---|---|---|
| RASGRF2 | -1.478816829 | 0.00239474 |
| SLCO4C1 | -1.753353931 | 0.01326495 |
| TREML2 | -1.889900259 | 0.00124586 |
| RHOU | -2.184386362 | 0.00255885 |
| B3GAT1 | -2.484371317 | 0.01326495 |
| GLI2 | -2.496629837 | 0.00189913 |
| PLXDC1 | -2.685498893 | 0.03136187 |
| RTL5 | -2.762275996 | 0.04009745 |
| KCNJ13 | -3.765415279 | 0.004636 |

common genes upregulated_NT vs CD28 & NT vs 41BB

| | |
|---|---|
| SLC22A17 | LURAP1L |
| ITGAD | SLC22A17 |
| ADCY6 | EPB41L4B |
| DAPK1 | ASAP2 |
| SAMD14 | TIE1 |
| PCGF2 | COL6A2 |
| KRT86 | MOXD1 |
| KRT81 | MMP9 |
| RUSC2 | MB |
| PALLD | COL6A1 |
| KRT7 | PTPN3 |
| HSPB1 | HMSD |
| SPINK2 | LHFPL6 |
| LINC01229 | PAWR |
| ADAMTS10 | KRT87P |
| HRH2 | ADCY6 |
| PITPNM2 | NLGN4Y |
| BTLA | EBI3 |
| FOXD1 | TLCD5 |
| AXIN2 | CTTNBP2NL |
| ME3 | ITGAD |
| PTGIS | SOX4 |
| BAIAP3 | KRT7 |
| CDCP1 | SOX18 |
| PLPP1 | KRT86 |
| PDE7B | PTGS2 |
| GSTA4 | RUSC2 |
| SPNS2 | NTRK1 |
| SIRPG | P2RY14 |
| NMUR1 | PLA2G4A |

common genes downregulated_NT vs CD28 & NT vs 41BB

| | |
|---|---|
| LPGAT1 | TC2N |
| NETO2 | TRIB2 |
| IFNG | NPC2 |
| RHOU | SAMD3 |
| GZMH | HS3ST3B1 |
| LUCAT1 | RASGRF2 |
| PLXDC2 | SLCO4C1 |
| SLC2A6 | TREML2 |
| | RHOU |
| | B3GAT1 |
| | GLI2 |
| | PLXDC1 |
| | RTL5 |
| | KCNJ13 |

| | |
|---|---|
| SOGA1 | SAMD14 |
| CDH24 | DAPK1 |
| H1-0 | NR5A2 |
| BACE2 | NBL1 |
| SERPINB6 | RTN4RL1 |
| ASPH | GPR35 |
| CLIP2 | PCGF2 |
| TPCN1 | KRT81 |
| HDAC5 | PALLD |
| FCMR | ICOSLG |
| MCOLN2 | EVC |
| | MSC |
| | MSC-AS1 |
| | FBN1 |
| | PANX2 |
| | EPB41L2 |
| | GATA3-AS1 |
| | C20orf204 |
| | BIRC3 |
| | ASB2 |
| | DUSP10 |
| | DLG4 |
| | HSPB1 |
| | HIC1 |
| | DNPH1 |
| | CHCHD10 |
| | SERTAD2 |

| CD28 vs 41BB | log2FoldChange | padj |
|---|---|---|
| TRBV11-3 | 3.65192637 | 0.00089172 |
| PLXND1 | 2.07940293 | 0.01249831 |
| PITPNM2 | 1.98332831 | 0.04491362 |
| PTGDR | 1.55444899 | 0.01460499 |
| FCMR | 1.0850748 | 0.00089172 |
| RCBTB2 | 0.80917634 | 0.02309391 |
| DNPH1 | -0.8450666 | 0.04246139 |
| NBL1 | -1.3368118 | 0.01460499 |
| BIRC3 | -1.4301873 | 0.00089172 |
| IFNG | -1.7649617 | 0.00018866 |
| SLC2A6 | -1.8693961 | 0.01101105 |
| MSC-AS1 | -2.1904003 | 0.02240241 |
| EGFL6 | -2.1922325 | 0.00013543 |
| TUBA5P | -2.2856035 | 0.01460499 |
| MSC | -2.7561813 | 0.00317726 |
| LUCAT1 | -2.7629678 | 0.00089172 |
| NLGN4Y | -3.2040866 | 0.04491362 |
| HMSD | -3.8700702 | 0.01460499 |
| COL6A2 | -3.9534839 | 0.01460499 |
| CLEC7A | -4.7058628 | 0.02986126 |
| MMP9 | -5.2321355 | 3.87E-10 |
| MB | -5.4809429 | 0.01212771 |
| COL6A1 | -5.877642 | 8.69E-13 |
| TIE1 | -6.9666444 | 0.00089172 |
